# Human blood vessel organoids reveal a critical role for CTGF in maintaining microvascular integrity

**DOI:** 10.1101/2022.09.01.505804

**Authors:** Sara G Romeo, Ilaria Secco, Edoardo Schneider, Christina M Reumiller, Celio XC Santos, Aman Pooni, Xiaoke Yin, Konstantinos Theofilatos, Silvia Cellone Trevelin, Lingfang Zeng, Giovanni E Mann, Andriana Margariti, Manuel Mayr, Ajay M Shah, Mauro Giacca, Anna Zampetaki

## Abstract

The microvasculature plays a key role in tissue perfusion, transport of mediators, and exchange of gases and metabolites to and from tissues. Microvascular dysfunction has emerged as an important contributor to cardiovascular diseases. In this study we used human blood vessel organoids (BVOs) as a model of the microvasculature to delineate the mechanisms of microvascular dysfunction caused by metabolic rewiring. BVOs fully recapitulated key features of the normal human microvasculature, including reliance of mature endothelial cells (ECs) on glycolytic metabolism, as concluded from metabolic flux assays using ^13^C-glucose labelling and mass spectrometry-based metabolomics. Treatment of BVOs with PFK15, a pharmacological inhibitor of glycolysis, resulted in rapid tissue restructuring, vessel regression with reduced pericyte coverage and alterations in tight junction morphology. Proteomic analysis of the BVO secretome revealed remodelling of the extracellular matrix and differential expression of paracrine mediators such as CTGF. Treatment with recombinant CTGF recovered tight junction formation and increased pericyte coverage in microvessels. Our metabolic and proteomics findings demonstrate that BVOs rapidly undergo restructuring in response to metabolic changes and identify CTGF as a critical paracrine regulator of microvascular integrity.

## Introduction

Microvascular dysfunction has emerged as a key contributor to cardiovascular diseases. Diabetic microangiopathy is the best characterised microvascular complication of diabetes mellitus^1, 2^, with the duration and severity of hyperglycemia being important determinants of vascular injury. Recently, the role of functional impairment of the microvasculature has been highlighted as a pathophysiological driver in heart failure^3^. A metanalysis of seventy-nine studies with a total of ∼60,000 individuals concluded that the direct readout of coronary microvascular dysfunction, namely the coronary flow reserve, is strongly associated with increased risk of all-cause mortality and major adverse cardiovascular events across a wide range of pathological processes. Therefore, a diagnostic and prognostic tool of microvascular dysfunction that could facilitate a more in-depth understanding of the disease, could also inform more precise patient selection for personalized novel therapeutic approaches^4, 5^.

The microvasculature mediates crucial functions, supplying oxygen and nutrients to tissues, controlling inflammation and exchanging gases and metabolites to and from tissues^6, 7^. Microvessels consisting of endothelial cells (ECs) and pericytes (PCs) form intricate networks in the microvasculature. These two cell types are in direct contact and deposit extracellular matrix (ECM) that forms the basement membrane, with PCs usually fully enclosed in this matrix. Disruption of the EC-PC crosstalk and PC loss from the microvasculature are hallmarks of microangiopathy^8^. Structural remodelling, including changes in vessel density, diameter and length along with alterations of capillary basal membrane thickness, impairs vessel function and can result in tissue hypoxia. In addition, dysregulated PCs display mechanical stiffness and altered contractility that contributes to abnormal EC behaviour, microvascular rarefaction and instability leading to tissue ischaemia^3, 8^. Although therapeutic strategies to enhance capillary sprouting and PC lining could stabilize the microvasculature and reduce tissue ischemia^9, 10^, the molecular mechanisms involved in EC and PC dysfunction are not well understood.

Recently, a human tissue model of the microvasculature has been developed, based on the generation of human blood vessel organoids (BVOs) from induced pluripotent stem (iPS) cells^11^. These self-organizing 3D tissue units can be grown in a Petri dish and display the morphological and molecular features of human microvasculature forming capillaries with a lumen, CD31^+^ endothelial lining, platelet derived growth factor receptor beta (PDGFRβ)^+^ pericyte coverage and the presence of a basement membrane^11^. BVOs derived from human iPS cells are particularly attractive for the development of personalized treatment strategies as they can faithfully recapitulate the human disease features and reflect the genetic heterogeneity of the human population. Thus, iPS-derived BVOs can serve as an experimental system to study the pathogenesis of microvascular dysfunction and as a system for drug and small compound screening^12^.

Metabolism has emerged as a key driver of angiogenesis in parallel to well-established growth factor signaling. In the microvasculature both ECs and PCs heavily rely on glycolysis for their energy demands^13, 14^. Phosphofructokinase-2/fructose-2,6-bisphosphatase 3 (PFKFB3) is a potent stimulator of glycolysis^15^. In mice genetic manipulation of PFKFB3 and pharmacological inhibition of PFKFB3 in *in vitro* experiments resulted in alterations in vascular sprouting and vessel formation^15^, indicating a key role for PFKFB3-driven glycolysis in the angiogenic potential.

Here, we use BVOs to investigate the mechanisms of metabolic rewiring-induced microvascular dysfunction and characterise the response of human microvessels to PFKFB3-driven glycolysis inhibition. Our experiments revealed that CTGF is a critical regulator of microvascular integrity, which could serve as a target for novel therapeutic interventions.

## Materials and Methods

### Cell culture

HEK293T cells were purchased from ThermoFisher Scientific and maintained in DMEM supplemented with 10% FBS. Cell culture images were taken using a Nikon Eclipse TS100 microscope and Nikon DS-Fil camera.

### PFK15 Treatment

The chemical selective 6-phosphofructo-2-kinase 3 (PFKFB3) inhibitor with the commercial name PFK15^16^ was purchased from Selleckchem. The inhibitor can be stored as stock solution of 50 mM in dimethyl sulfoxide (DMSO) at −80°C. Once thawed, aliquots may be kept at 4°C for 2 weeks. Unless otherwise stated, BVOs or iPS-ECs were washed twice and then incubated in EBM2 with no supplement (BVOs for 2h, iPS-ECs for 1h). Subsequently, they were treated for the indicated time with PFK15 (2.5µM) or DMSO in EBM2 media in the absence of supplements.

### Generation of Blood Vessel Organoids

BVOs were generated from iPS cells using the protocol established by Wimmer et al, 2019^17^. The KOLF2 human iPS cell line was obtained from the Wellcome Trust Sanger Institute. hiPSCs were plated on Matrigel-coated (ATCC, ACS-3035) 6 well plates and cultured using the StemMACS iPS Brew-XF system (Milteney Biotech, 130-104-368). For differentiation to BVOs, 2.5×10^5^ iPSCs per well were seeded into Ultra-low adherent 6-well plates (Appleton Woods, 3471) in Aggregation Media to form cell aggregates (diameter average 50-150 µm) for approximately one day. Aggregates were collected by gravitation (Day 0) and resuspended into the mesodermal induction media (N2B27 Media supplemented with 12 μM CHIR (Tocris Bioscience, cat. no. 4423) and 30 ng/ml BMP4 (ThermoFisher, PHC9534)^17^. On day 3, the aggregates were collected by gravitation and plated in N2B27 media supplemented with 100ng/ml VEGFA (Peprotech, 100-20) and 2 μM Forskolin (R&D system, 1099) for vascular lineage promotion. On day 5, the aggregates were embedded in a substrate of Collagen I-Matrigel (Advanced BioMatrix, 5005) (ratio 4:1) with StemPro-34 SFM (Gibco, 10639011) supplemented with 15% FBS (Thermo, 10500064), 100ng/ml VEGFA and 100ng/ml FGF2 (Milteney Biotech, 130-093-841) to induce vascular network formation^17^. The sprouting appeared within two days in all preparations. Fresh media was provided after three days and then every other day. Vascular networks (VNs) were established between days 10 to 12 and extracted from the Matrigel to a 96-well ultra-low-attachment plate (2BScientific, MS-9096UZ) where they self-assembled to BVOs. BVOs were kept in culture for up to 40 days.

### Differentiation of iPS-EC

The method used was adapted from previously published protocols^18^. iPS cells were seeded on Corning Matrigel Growth Factor Reduced (GFR) Basement Membrane Matrix (SLS, 356231) at a density of 1.6×10^5^ cells per 6 well in StemMACS iPS-Brew XF (Milteney Biotech, 130-104-368) supplemented with 10 μM ROCK inhibitor Y-27632 (ATCC ACS-3030) for 24 h. The following day (day 1), the medium was changed to N2B27 medium, a 1:1 ratio of Neurobasal medium (Thermo, 21103049) and DMEM/F12 (Gibco, 11330-032), with N2 (Thermo, 17502048), B27 (Thermo, 12587010), Glutamax (Thermo, 350050061) and freshly supplemented with 8 μM of CHIR (Sigma, SML1046) and 25 ng/ml of BMP4 (ThermoFisher, PHC9534). After 72h (day 4), the medium was replaced with StemPro-34 SFM (Gibco, 10639011) supplemented with 200 ng/ml human VEGFA (Peprotech, 100-20) and 2 μM Forskolin (R&D system, 1099). On day 6, cells were selected by Magnetic Activated Cell Sorting (MACS) for iPS-EC expressing CD144 using Microbeads Kit (Miltenyi, 130-097-857). Positive cells were seeded on mouse Collagen IV (Biotechne, 3410-010-02) coated plate in EGM2-MV medium (Promocell, C-22011) supplemented with 20% FBS (Thermo, 10500064) and 50 ng/ml VEGFA. Cells were used for experiments up to 3 passages.

### Metabolic Analyses

To assess the metabolic fate of glucose, iPS-ECs were cultured with 5mM U-^13^C_6_-glucose (Cambridge Isotope Laboratories) in DMEM without glucose, glutamine, and pyruvate (ThermoFisher Scientific) supplemented with 1% dialyzed FBS, 1% penicillin-streptomycin, and 1% L-glutamine for 7 h, as pilot studies indicated that cells reached isotopic steady-state by this time. The medium was then removed and 80:20 methanol: water (−80 °C, extraction solvent) was added to cells for 15 min at − 80 °C for metabolic quenching. Cells were scraped in the extraction solvent and cell lysates pipetted into Eppendorf tubes followed by a sonication step. Then, samples were centrifuged at 16,000 g for 10 min at 4 °C to pellet debris.

The supernatant was transferred to a new tube and dried using a SpeedVac (Thermofisher Scientific, Savant, SPD131DDA). The dried pellet was resuspended in 50 µl chloroform /100 µl methanol/100 µl H_2_O. The top, polar fraction was further dried and stored at −80 °C until LC-MS analysis. All metabolite analyses were performed on three biological replicates.

### LC-MS analysis

The LC-MS method has been previously^19^. Briefly, the dried polar metabolite fractions were reconstituted in acetonitrile/water (v/v 3/2), vortexed and centrifuged at 16,000 g for 3 min before analysis on a 1290 Infinity II ultrahigh performance liquid chromatography (UHPLC) system coupled to a 6546-quadrupole time-of-flight (Q-TOF) mass spectrometer (Agilent Technologies). Samples were separated on a Poroshell 120 HILIC-Z column (100 × 2.1 mm, 2.7 μm, Agilent) attached to a guard column (5 × 2.1 mm, 2.7 µm) and analysed in negative ionization mode using water with 10mM ammonium acetate (solvent A) and acetonitrile with 10mM ammonium acetate (solvent B), both solvents containing 5 µM Infinity Lab deactivator additive (Agilent Technologies). The elution gradient used was as follows: isocratic step at 95% B for 2 min, 95 to 65% B in 12 min, maintained at 65% B for 3 min, then returned to initial conditions over 1 min, and then the column was equilibrated at initial conditions for 8 min. The flow rate was 0.25 mL/min; the injection volume was 1μL, and the column oven was maintained at 30 °C. Feature annotation and metabolite identification were based on accurate mass and standard retention times with a tolerance of ±5 ppm and ±0.5 min, respectively, and performed with MassHunter Profinder (version 10.0.2, Agilent Technologies) using our in-house curated metabolite library based on metabolite standards (Sigma-Aldrich). C^13^ label incorporation levels were normalised to the natural occurrence of C^13^ isotopes and are represented as corrected abundance percentages. Samples were run in one batch and injected into technical duplicates. Downstream pathway enrichment analysis was performed using Metaboanalyst 5.0 Software^20^.

### Immunohistochemistry

BVOs were rinsed twice in PBS and then fixed in 4% paraformaldehyde (PFA) for 1 h at room temperature (RT) and washed twice in PBS. BVOs were dehydrated in 20% sucrose at 4°C overnight, then embedded in Gelatin solution and stored at − 20°C for cryosectioning. The frozen BVOs were sectioned using NX70 Cryostat (Thermo Scientific) to 20 and/or 70-μm thickness. Frozen sections were then washed in PBS for 5 minutes and then permeabilized and blocked in Blocking buffer for 2h at RT. Primary antibodies were diluted in Blocking buffer and incubated overnight in a cold room. Antibodies used and dilution factors’ information are available in Supplementary Table S1. On the following day, sections were washed three times in PBS with 0.1% TritonX-100 (PBS-T) and then incubated with labelled secondary antibodies for 2 h at RT. After two washes in PBS-T the sections were counterstained with DAPI and mounted with Fluoromount-G mounting media (Thermo, 00-4958-02).

For immunostaining intact BVOs and VNs were rinsed in PBS then fixed in 4% PFA for 30 minutes at RT and washed twice in PBS. The fixed VNs were stored in PBS at 4°C for up to a month. Stained BVOs were mounted into iSpacer (0.5 mm deep imaging Spacer, SunJin Lab). Samples were viewed and imaged using the Spinning Disk Confocal System (Nikon) and the Operetta CLS High-Content Analysis System (PerkinElmer) at a 20x magnification.

### Quantitative Reverse Transcription PCR (qRT-PCR)

RNA was isolated using the PureLink RNA Mini kit (ThermoFisher Scientific, 12183018A) according to the manufacturer’s recommendation and reverse-transcribed into cDNA using High-Capacity cDNA Reverse Transcription Kit (ThermoFisher, 4368813). The primer sequences are provided in Supplementary Table S2. The reactions were conducted using SYBR green master mix (ThermoFisher, 4472920) on the ViiA 7 real-time PCR system (ThermoFisher) at 95 °C for 10 min, followed by 40 cycles of 95 °C for 15 sec and 60 °C for 1 min. Beta-actin was used as a normalization control.

### Flow Cytometric Analysis

BVOs (n=7 per group) were mechanically dissociated using a scalpel and then incubated in Dissociation solution (1.7mg Dispase, 0.2mg Liberase and 0.1mg DNase per ml) in PBS for 20 min at 37°C. BVO solutions were passed up to 10 times through 21g needles. Approximately 50,000 single cells were resuspended in 100μl of FACS buffer (PBS containing 1% FBS) and stained with fluorescence-conjugated antibodies for 30 min at 4°C in the dark. Information for the antibodies used and dilution factors is listed in Supplementary Table S3. The cells were washed in PBS and resuspended in 1% PFA in PBS. Data were acquired the following day using a Fortessa Flow Cytometer analyzer (BD) and analyzed using FlowJo software (Becton & Dickinson and Company).

### Western blot

Whole protein lysates were isolated from iPS cells and iPS-ECs using the 1x Cell Lysis buffer (Cell Signalling, 9803) supplemented with protease inhibitors (Roche 11697498001). Cells were washed using PBS and harvested at the same passage and confluency (80%) to minimize differences between samples. Proteins were quantified using the BCA method (ThermoFisher Scientific, 23227). A total of 16 μg of proteins were loaded and separated on 4-12% polyacrylamide-SDS gels under reducing conditions (Invitrogen, NP0322) and transferred to nitrocellulose membranes (Amersham, 10600003).

Membranes were blocked in 0.1% Tween20 TBS (TBS-T) and 5% dry non-fat milk for 1 h at RT and incubated with the different primary antibodies overnight at 4°C. The following day, membranes were washed three times for 10 min with TBS-T and incubated for 1.5 h at RT with the respective HRP (horseradish peroxidase)-conjugated secondary antibodies (anti-rabbit, 211-032-171, and anti-mouse, 115-035-174, Stratech with 1:4000 dilution). The ECL detection system (GE Health GERPN2232) was used. The antibodies used and dilution factors’ information are listed in Supplementary Table S4.

### Zymography

The conditioned media was harvested from BVOs following 24h of treatment and cell debris was removed by centrifugation at 4000 rpm for 10 min. The supernatant was transferred into a new tube and stored at −80°C. Samples were mixed with non-reducing Tris-Glycine SDS sample buffer (2x, ThermoFisher) and loaded to a Novex® 10% zymogram gel (gelatin, ThermoFisher). After electrophoresis, the gel was renatured and developed at 37°C overnight according to the manufacturer’s recommendations. The gel was stained with Simply blue Safestain (ThermoFisher).

### Luciferase reporter assays

Luciferase assays were performed in HEK293T cells using the YAP/TAZ reporter 8xGTIIC-lux (Addgene, 34615). *Renilla*-Luc (0.05 µg/well) was included as an internal control. Cells were plated in 12 well plates (10^5^ cells/ 12w) and the following day transfected using Lipofectamine 2000 (ThermoFisher Scientific) transfection reagent. The PFK15 and recombinant CTGF as indicated per condition were supplemented to the wells two hours after the initiation of transfection. Luciferase activity was detected 24 h after transfection and normalized to Renilla luciferase activity using the Dual-Glo Luciferase system (Promega). The relative luciferase unit was defined as the ratio of Firefly versus Renilla. At least three independent transfections were performed in quadruplicates.

### *In vitro* tube formation assay

For *in vitro* tube formation, 24-well plates were coated with 200 μl/well of Matrigel Matrix (Corning). The plates were incubated at 37°C for 30 min. Subsequently, 6×10^4^ iPS-ECs were plated in each well and incubated for 4 hours, at 37°C in a humidified incubator supplemented with 5% CO_2_. Vascular network images were taken using a Nikon Eclipse TS100 microscope and Nikon DS-Fil camera.

### Proteomic Analysis of the BVO secretome

Secreted proteins were concentrated using 3kD MWCO spin filters (Amicon) at 20,000 x g, 4°C. The protein samples were denatured by 6M urea, 2M thiourea and reduced by 10mM DTT at 37°C for 1h, alkylated by 50mM iodoacetamide at RT in the dark for 1h, and washed with 0.1M triethylammonium bicarbonate (TEAB, pH=8.5) for 3 times in the spin filter. The proteins were digested by trypsin/lysC (protein: enzyme = 25:1, Millipore) at 37°C overnight. Peptides were acidified by 1% trifluoroacetic acid (TFA) and C18 cleanup was performed using Bravo AssayMAP robot with C18 cartridges (Agilent) following the manufacturer’s instruction. Eluted peptides were SpeedVac dried and resuspended in 2% acetonitrile (ACN), 0.05% TFA in LC-MS grade H_2_O.

Peptide samples were injected and separated by an UltiMate3000 RSLCnano system (EASY-Spray C18 reversed-phase column, 75µm × 50cm, 2 µm, Thermo Fisher Scientific) using the following LC gradient: 0-1 min: 1% B; 1-6 min: 1-6% B; 6-40 min: 6-18% B; 40-70 min: 18-35% B; 70-80 min: 35-45% B; 80-81min: 45-99% B; 81-89.8 min: 99% B; 90-120 min: 1% B (A=0.1% formic acid in H_2_O, B=80% ACN, 0.1% formic acid in H_2_O). The separated peptides were directly injected into an Orbitrap Q Exactive HF Mass Spectrometer (Thermo Fisher Scientific). Full MS spectra were collected using Orbitrap with scan range of 375-1500 m/z and a resolution of 60,000. The most abundant 15 ions from the full MS scan were selected for data-dependant MS2 with HCD fragmentation and acquired using Orbitrap with a resolution of 15,000 and isolation windows 2 m/z. Dynamic exclusion of 40 seconds and lock mass of 445.12003 m/z were used.

RAW data were analysed using Proteome Discoverer (version 2.4, Thermo Fisher Scientific) with MASCOT algorithm (version 2.6.0, Matrix Science). The following parameters were used: UniProt/SwissProt human and bovine protein database (version 2021_01, 26410 protein entries) were used; trypsin was used as enzyme with maximum 2 missed cleavages allowed; carbamidomethylation on cysteines was selected as static modification and oxidation on methionine, lysine and proline was selected as dynamic modifications; precursor ion mass tolerance was set at 10ppm, and fragment mass tolerance was set at 20 milli mass unit (mmu). Protein identification FDR confidence was set to High and the minimum number of peptides per protein was 2. The precursor peak area was used for quantification and exported for further statistical analysis.

### Bioinformatic analysis

The proteomics data were first normalized using total ion intensity. Then bovine proteins or human proteins with a bovine homolog that had more matched spectra in the bovine protein compared to the human were filtered out. The dataset was further filtered to keep only proteins with less than 30% missing values. All remaining missing values were imputed with the KNN-Impute method with k equal to 2. The relative quantities of the proteins were scaled using log2 transformation.

The limma package^21^ has been used to compare between different phenotypes using the Ebayes algorithm and correcting for selected covariates. The nominal p-values were adjusted for multiple testing using Benjamini-Hochberg method^22^ but because of the limited sample size, a combination of thresholds of 0.05 on the nominal p-values and 0.25 for the absolute log2 fold change were used to infer statistically significant changes. Beanplots, Volcano Plots and heatmaps plots were constructed using the Beanplots ^23^,Ggplot2 ^24^ and Corrplot ^25^ packages of R programming environment^26^, using R version 4.2.1.

Network visualizations were conducted using Cytoscape tool ^27^ with protein-protein interaction networks being reconstructed from String web tool ^28^. Pathway and functional enrichment analysis was conducted using David tool ^29^. This analysis included pathway terms from Reactome data repository ^30^, KEGG ^31^ and functional terms from Gene Ontology ^32^. Significantly enriched terms were inferred with a Benjamini-Hochberg adjusted p-value threshold of 0.05. Transcriptional factor enrichment analysis was performed using the ChEA3 tool ^33^ using the ENCODE ChIP-sequencing data ^34^.

### Statistics

Biological replicates were performed in all the experiments with n = 3 independent preparations. Data are expressed as the mean ± SD and were analyzed using GraphPad Prism 9 software with a two-tailed Student’s t-test for two groups or ANOVA for more than two groups. A value of *p < 0.05, **p < 0.01, ***p < 0.001 was considered significant.

## Results

### Generation of human blood vessel organoids (BVOs)

To study the mechanisms of metabolic rewiring induced microangiopathy, we generated BVOs employing the protocol established by Wimmer et al^17^. Human iPS cells were used to generate cell aggregates with a diameter ranging from 50-150 μm. These cell aggregates subsequently went through mesoderm (days 0-3) and vascular lineage differentiation (days 4-5) and were then embedded in a 3D collagen I-Matrigel matrix in 12-well plates. Vascular networks (VNs) with clearly visible tip cells emerged within 2-3 days. After 5-7 days of sprouting in the 3D substrate, VNs were extracted and transferred into an ultra-low 96 well adhesion plate, where they self-assembled into BVOs (Figure 1a). BVOs consisting of ∼30,000 cells per organoid, with an average diameter 1.5-2mm (data not shown) were maintained in suspension culture in a low-attachment 96 well U bottom plate for the duration of the experiment. Immunofluorescence staining was used to characterize the VNs and BVOs and demonstrated the presence of a dense vascular sprouting comprised of CD31^+^ ECs, PDGFRβ^+^ mural cells and a thick deposition of basement membrane as indicated by the detection of Collagen IV (ColIV) in both VNs and whole BVOs (Figure 1b). A more detailed analysis of the VN structure revealed PDGFRβ^+^ mural cells tightly associated with ECs, with single PDGFRβ^+^ cells elongated along ECs and embracing the vessel wrapping several ECs incompletely (Figure 1c) suggesting that these cells are PCs (Figure 1d). Flow cytometry analysis of digests of BVOs further confirmed the presence of CD31^+^ ECs, PCs as PDGFRβ^+^ cells, mesenchymal stem-like cells as CD90^+^CD73^+^CD44^+^ cells and haematopoietic cells as CD45^+^ (Figure 1e). These characteristics indicate that iPS derived BVOs recapitulate key aspects of the human microvasculature and can be used as a model to study the mechanisms of microangiopathy.

**Figure 1.**
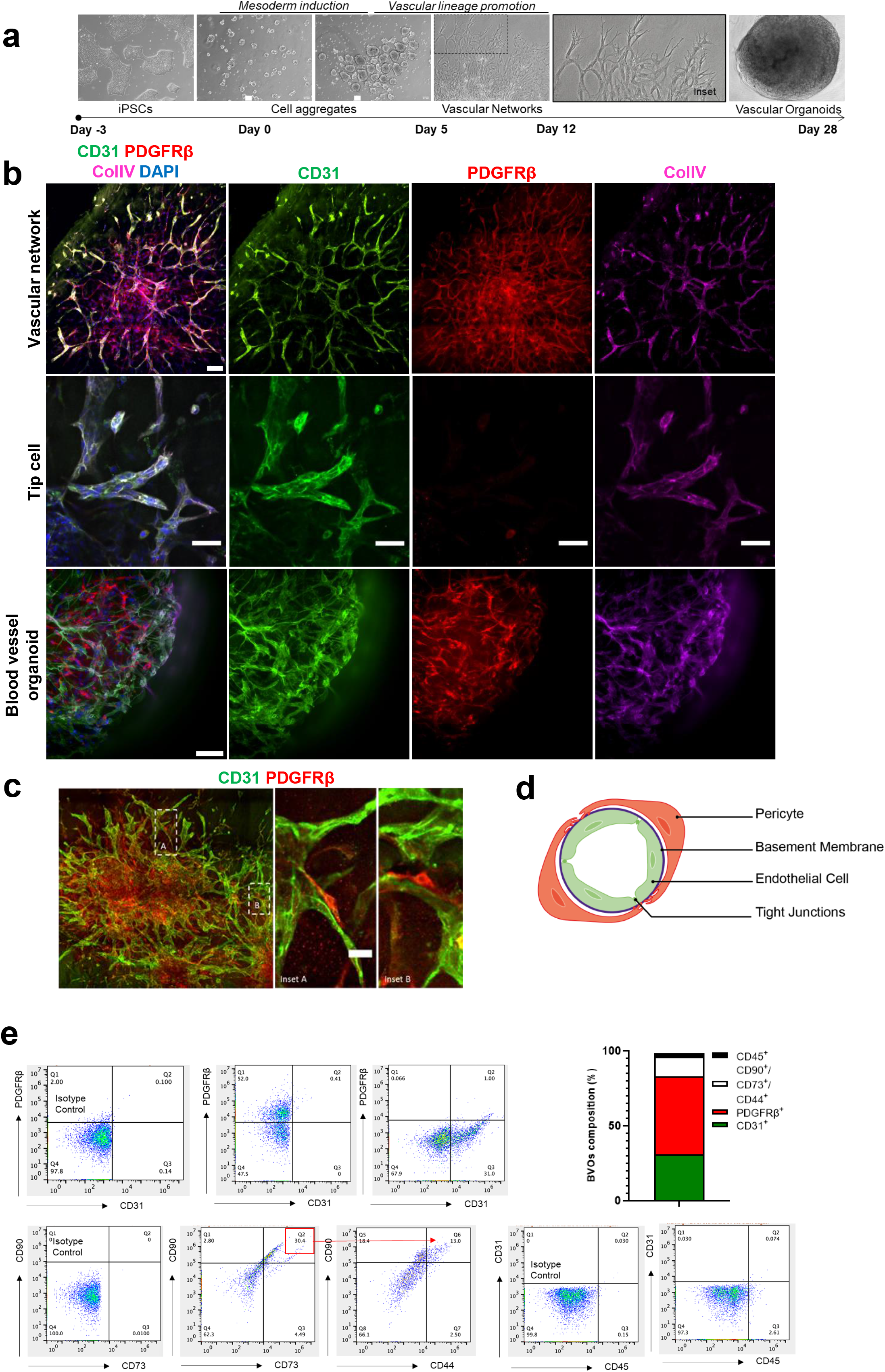
Generation of human blood vessel organoids (BVOs). **(a)** Bright-field images of human iPS cell differentiation into vascular networks and BVOs. **(b)** Phenotypic characterization of vascular networks and BVOs, showing CD31-expressing ECs (green) forming vascular networks covered by PDGFRβ^+^ pericytes (PCs, red) and a basement membrane (collagen type IV (Col IV), magenta). **(c)** Immunofluorescence confocal imaging of vascular networks showing pericyte coverage (PDGFRβ^+^, red) of CD31^+^ ECs (green). **(d)** Schematic depicting the microvascular structure. **(e)** FACS was used to determine the different cell populations in BVOs. ECs were defined as CD31^+^, PCs as PDGFRβ^+^, mesenchymal stem-like cells as CD90^+^CD73^+^CD44^+^ and haematopoietic cells as CD45^+^. Bar scales 100 μm and 50 μm.

### IPS derived ECs are highly glycolytic

Vessel sprouting and angiogenesis are directly linked to the metabolic profile of ECs^14^. Mature ECs rely on glycolysis rather than oxidative phosphorylation for energy production^15^. Aiming to determine whether ECs differentiated from iPS cells have a similar functional and metabolic profile as mature ECs, we generated iPS-derived ECs using a stepwise differentiation protocol to obtain mesodermal progenitors and subsequently vascular lineage cells^18^. CD144^+^ cells from this population were isolated using MACS and their angiogenic potential was tested in a tube formation assay (Figure 2a). Immunofluorescence staining confirmed successful differentiation, detecting pluripotency markers in iPS cells (Figure 2b) and the EC markers CD31, CD144 and tight junction protein 1 (ZO1) in the iPS-EC cells (Figure2c). These results were also confirmed by western blot analysis (Figure 2d).

**Figure 2.**
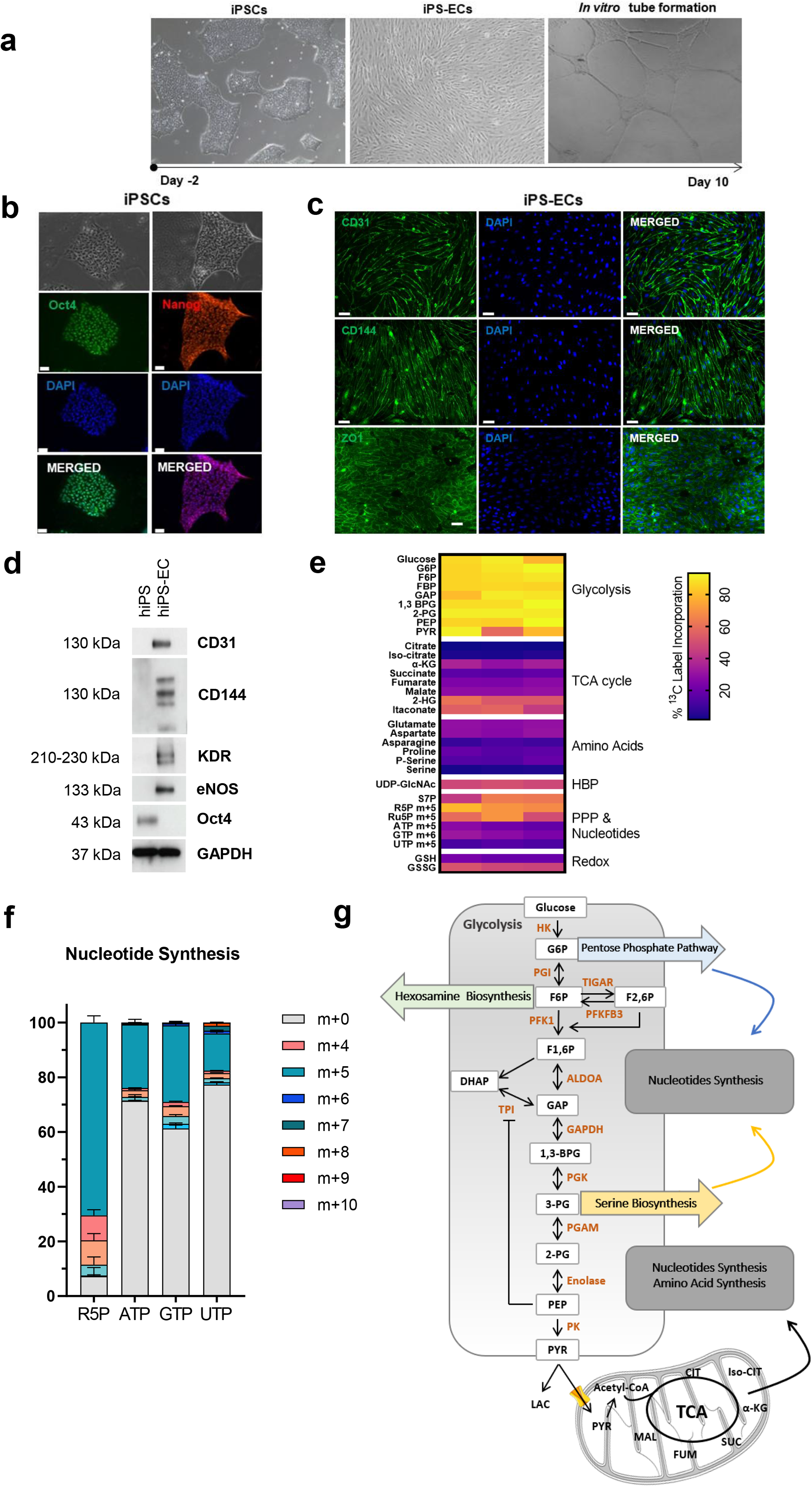
Human iPS cells differentiation toward ECs. **(a)** Typical morphology of a human iPS colony, iPS derived ECs (iPS-ECs) and iPS-EC *in vitro* tube formation indicating their angiogenic potential are shown by bright field microscopy. **(b)** Immunofluorescence confocal image demonstrating the expression of pluripotency markers in human iPS cells and **(c)** EC-specific markers CD31, CD144 and ZO1 in iPS-ECs. **(d)** The expression of EC markers was confirmed by Western blot. **(e)** Heat map illustrating ^13^C label incorporation determined by stable isotope tracing of U-^13^C_6_ glucose. **(f)** Mass isotopologue distribution of nucleotides and metabolites related to de-novo nucleotide synthesis. **(g)** Schematic representation of glycolysis with highlighted metabolic pathways that contribute to biomass production. Abbreviations: **HK** (hexokinase), **G6P** (glucose-6-phosphate), **PGI** (phosphoglucose isomerase), **F6P** (fructose-6-phosphate), **TIGAR (**TP53-induced glycolysis and apoptosis regulator), **PFKFB3 (**6-phosphofructo-2-kinase/Fructose-2,6-biphosphatase 3), **F-2**,**6-BP (**fructose-2,6-bisphosphate), **PFK1 (**phosphofructokinase-1), **F1**,**6 BP (**fructose-1,6-bisphosphate), **ALDOA (**aldolase A), **GAP (**glyceraldehyde 3-phosphate), **GAPDH** (glyceraldehyde-3-phosphate dehydrogenase), **DHAP (**dihydroxyacetone phosphate), **TPI (**triosephosphate isomerase), **1**,**3-BPG (**1,3-bisphosphoglycerate), **PGK** (Phosphoglycerate kinase), **3-PG (**3-phosphoglycerate), **PGAM (**phosphoglycerate mutase), **2-PG (**2-phosphoglycerate), **PEP (**phosphoenolpyruvate), **PK (**pyruvate kinase), **HBP** (Hexosamine biosynthetic pathway), **PPP** (Pentose phosphate pathway), **R5P** (Ribose 5-phosphate), **S7P** (Sedoheptulose 7-phosphate), **UDP-GlcNAc** (Uridine diphosphate N-acetylglucosamine), **P-Serine** (Phosphoserine), α **-KG** (α-Ketoglutaric acid), **2-HG** (2-Hydroxyglutarate). Data from n = 3 independent experiments. Bar scales 50 μm.

Endothelial metabolism has emerged as a major determinant of the angiogenic process^14^. Hence, we evaluated the metabolic profile of iPS-ECs. Targeted stable isotope tracing of U-^13^C_6_ glucose demonstrated that iPS-ECs heavily rely on glucose for biomass production (Figure 2e). In particular, iPS-ECs show high levels of ^13^C label incorporation from glucose into nucleotides (ATP, GTP, UTP) and metabolites related to de-novo nucleotide synthesis, *i*.*e*. the pyrimidine precursor ribose-5 phosphate (compare Figure 2f). These data demonstrate that iPS cells differentiated to ECs display key morphological, functional properties of mature ECs and a metabolic profile that indicates a strong reliance on glycolysis for energy production.

### Inhibition of glycolysis alters the BVOs cellular composition

Glycolysis in ECs is regulated by PFKFB3, a phosphofructokinase-2/fructose-2,6-bisphosphatase enzyme that synthesizes fructose-2,6-bisphosphate (F2,6P2), an allosteric activator of PFK-1, the most potent stimulator of glycolysis (Figure 2g)^15^. Genetic targeting of PFKFB3 in mice and pharmacological inhibition in *in vitro* experiments results in alterations in vascular sprouting and vessel formation^15^, indicating a key role for PFKFB3-driven glycolysis in the angiogenic process.

In human iPS derived cells, we found that PFK15^16^, a chemical inhibitor of PFKFB3 does not affect cell survival of iPS-ECs up to a concentration of 5μM (Supplementary Figure 1). Aiming to go beyond the response of a single cell type and delineate the impact of a short-term PFKFB3 inhibition in the human microvascular tissue, we employed BVOs for further studies. BVOs represent a human model of self-assembled 3D microvascular tissue that can offer unique insights into the mechanisms of tissue remodelling. It can overcome interspecies variations and is more physiologically relevant to coculture systems. Additionally, BVOs can be used to determine cell to cell and cell to ECM interactions, hence providing a better model of the human microvasculature^12, 35^.

We found that after a 24h PFK15 treatment no modification in BVO diameter or organoid disintegration was observed (Figure 3a, 3b). This is important as changes in organoids size have been previously linked to their function ^36^. Although no significant difference in cell numbers in the treated and untreated organoids was detected (Figure 3c), inhibition of glycolysis led to a significant decrease in proliferating cells (Figure 3d) accompanied by an increase in apoptosis (Figure 3e). Further analysis indicated that the EC CD31^+^ fraction displayed reduced proliferation (Figure 3f), but no difference in apoptosis (Figure 3g). No statistically significant differences in proliferation or apoptosis were observed in the PDGFRβ^+^ cells (Figure 3h, 3i). These findings indicate that inhibiting PFKFB3-driven glycolysis in BVOs leads to alteration in the cellular state, but not to organoid disintegration.

**Figure 3.**
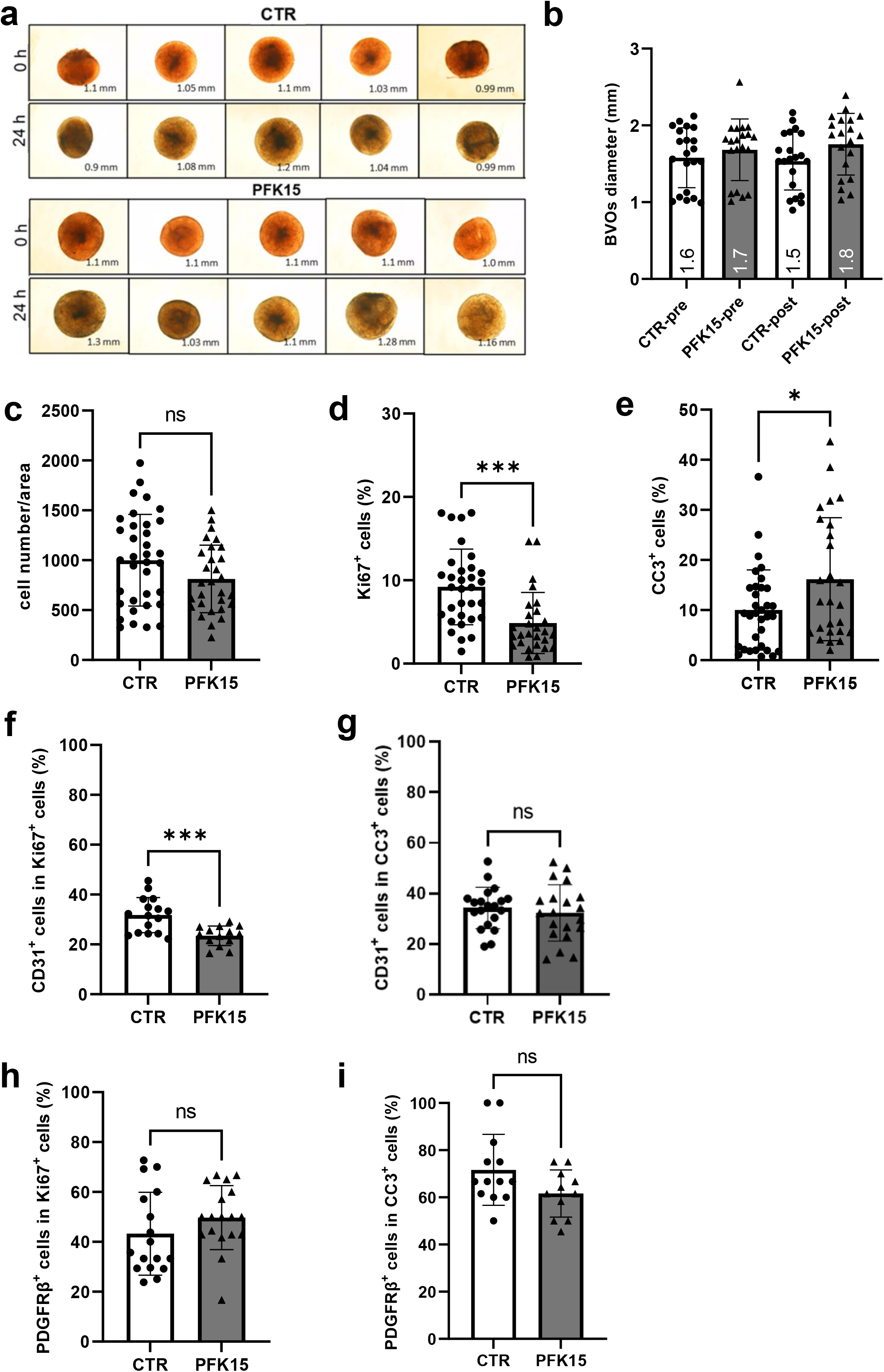
The effect of PFK15 on BVO diameter and cellular composition. **(a)** Bright-field images of BVOs treated with DMSO (CTR) or PFK15 (2.5µM) for 24h and **(b)** quantification of BVO diameter pre- and post-treatment n=20 BVOs per group. **(c)** Total number of cells per area, **(d)** percentage of proliferating cells and **(e)** percentage of cleaved caspase 3 (CC3) positive cells. N=28 BVOs per group **(f, g)** CD31^+^ proliferating cells and CD31^+^ cleaved caspase 3 positive cells respectively. N=16 per group **(h, i)** PDGFRβ^+^ proliferating cells and PDGFRβ^+^ cleaved caspase 3 positive cells respectively. N=12 per group. Values are presented as mean ± SD; P values were calculated using a two-tailed Student’s t-test. (* <0.05; **<0.005; ***<0.0005). Bar scales 200 μm. P values were corrected for multiple testing using Original FDR Benjamini-Hochberg method; q-values are provided in Supplementary Table 5.

### Inhibition of glycolysis triggers structural remodelling of BVOs and reduced pericyte coverage

BVOs display striking similarity with the human microvasculature, consisting of a dense network of capillaries with CD31^+^ endothelial microvessels with PDGFRβ^+^, chondroitin sulfate proteoglycan 4 (NG2)^+^ mural cells and pericyte coverage of microvessels (Figure 4a, Supplementary Figure 2, supplementary movies 1, 2). Interestingly, even a short 24h inhibition of PFKFB3-driven glycolysis led to structural modification of BVOs and resulted in reduced pericyte coverage by>50% (Figure 4a, 4b). In PFK15 treated BVOs, pericyte dropout was observed with a significant portion of PDGFRβ^+^ cells becoming ‘extravascular’ and not attached to CD31^+^ ECs^37^. Additionally, quantification of microvessels revealed that in PFK15 treated BVOs, the vessel density and the vessel length (Figure 4c, 4d and Figure 4e respectively) decreased significantly. Likewise, the vessel diameter in PFK15 treated BVOs was reduced, indicating vessel constriction (Figure 4f). These findings demonstrate that in response to alteration in glycolysis BVOs undergo rapid structural remodelling and microvessel regression.

**Figure 4.**
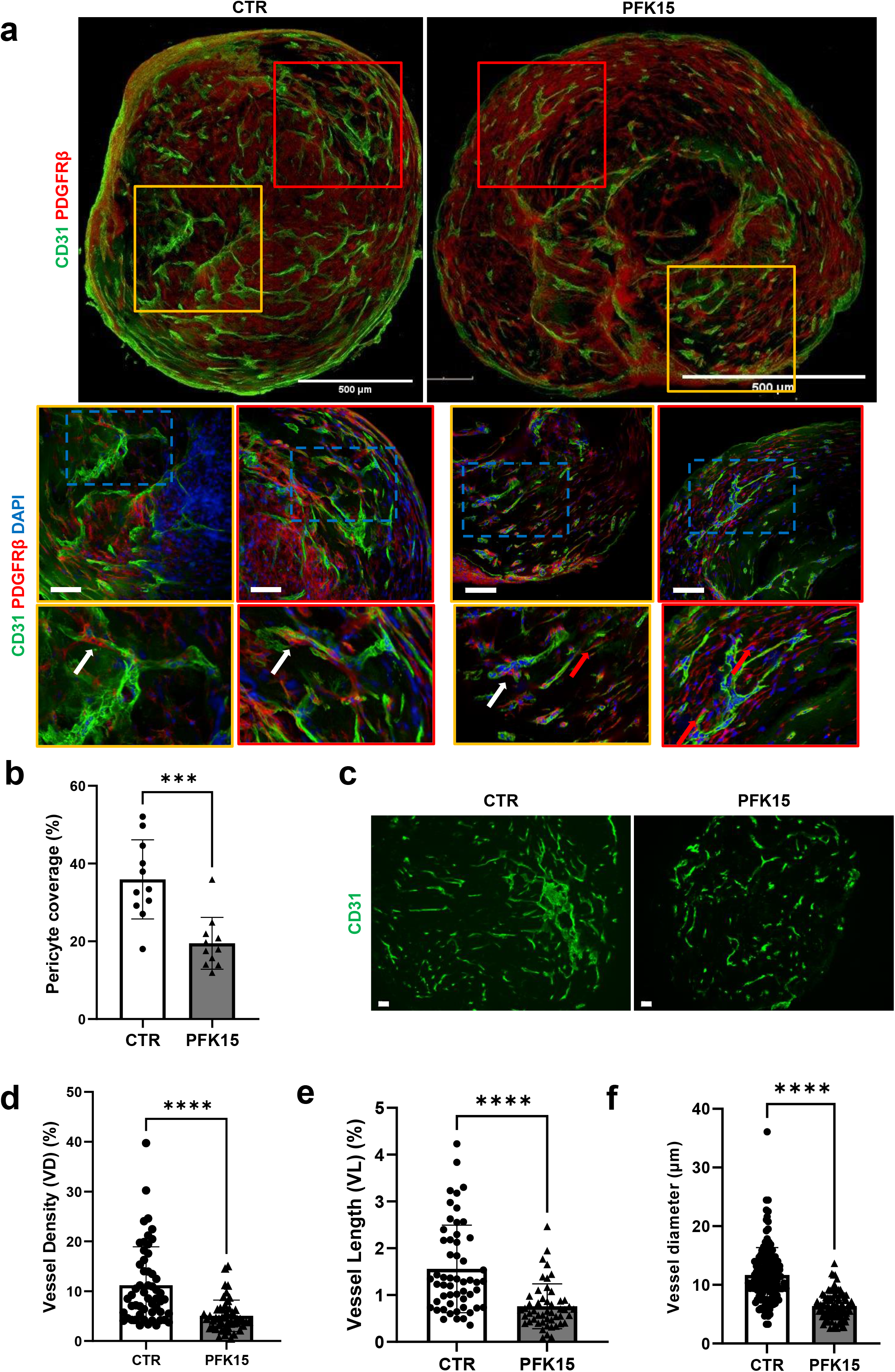
The effect of PFK15 on BVO structure. **(a)** Immunofluorescence confocal imaging of BVO sections showing pericyte coverage (PDGFRβ, red) of CD31^+^ ECs (green). White arrows indicate pericytes attached to microvessels, red arrows indicate ‘extravascular pericytes’. **(b)** Percentage of pericyte coverage n=2 BVOs per group, 3 separate preparations. **(c)** Differential vessel density as detected by immunofluorescence imaging of CD31^+^ ECs (green) on BVO sections. **(d)** Quantification of vessel density, **(e)** length in n=2 BVOs per group, 3 separate preparations in 4 different areas per x10 images and **(f)** vessel diameter in n=100 BVOs’ vessels (total cross-sections analysed from 3 separate preparations per condition in 70μm sections). Values are presented as mean with SD; P values were calculated using a two-tailed Student’s t-test. (* <0.05; **<0.005; ***<0.0005; ****<0.0001). Bar scales 50 and 500 μm.

### Pericyte differentiation to smooth muscle-like cells in BVOs following inhibition of glycolysis

Signals derived from the microvascular endothelium can regulate pericyte growth and differentiation. However the effect of metabolic reprogramming in the PC differentiation state is unknown^38, 39^. To determine the effect of inhibition of glycolysis in the PC phenotype we evaluated the expression of α-smooth muscle actin (αSMA), a gene conferring contractility and a commonly used marker of smooth muscle cells (SMCs). Immunofluorescence staining of BVOs revealed a 2.5-fold increase in the αSMA positive area in PFK15 treated BVOs (Figure 5a, 5b), with more than 60% of PDGFRβ^+^ cells also expressing αSMA (Figure 5c). Similarly, a significant but less pronounced increase in transgelin (SM22), another marker of SMCs was observed in the PFK15 treated BVOs (Figure 5b, 5c). Intriguingly, qPCR assessment did not reveal an upregulation of αSMA gene expression in BVOs after 24h of PFK15 treatment (Figure 5d). Previous studies have shown that TGFβ1 stimulation in retinal pericytes cultured *in vitro* triggers an early αSMA mRNA upregulation at 4h, with levels declining to baseline after 14h hours ^40^. Therefore an earlier time point was assessed. Even after 4h of PFK15 treatment no increase in the levels of αSMA mRNA was detected by qPCR (Figure 5e), indicating that the observed αSMA increase does not involve *de novo* protein synthesis but rather posttranscriptional regulation. These data demonstrate that inhibition of glycolysis triggers pericyte dropout from the microvessels and increased pericyte differentiation to SMC-like cells.

**Figure 5.**
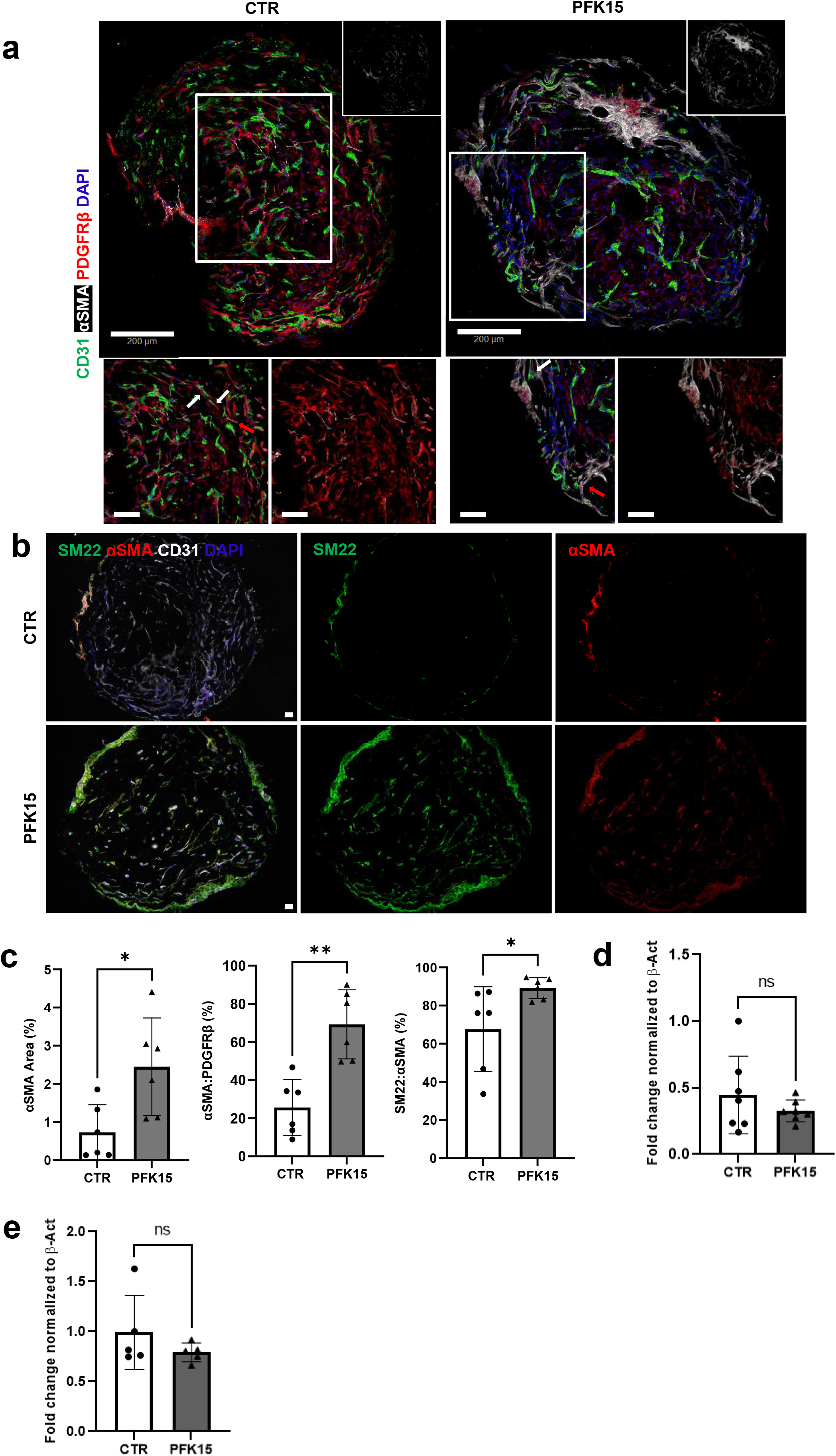
The effect of PFK15 on PC differentiation to Smooth Muscle-like cells in BVOs. **(a)** Phenotypic characterization of alpha smooth muscle actin (αSMA, white) expression in BVO sections using immunofluorescence confocal microscopy. PDGFRβ^+^ cells are shown in red and CD31^+^ ECs in green. **(b)** Phenotypic characterization of SM22 (green) and αSMA (red) expression in BVO sections using immunofluorescence confocal microscopy. CD31^+^ ECs are shown in white. **(c)** quantification of αSMA area, αSMA and PDGFRβ^+^ area and αSMA and SM22 area in BVOs following treatment with DMSO (CTR) or PFK15 (2.5µM) for 24h. n=2 BVOs, 3 separate preparations. (**d**) αSMA gene expression as assessed by qPCR after 24h of PFK15 treatment. β-actin was used as a normalization control. n=7 BVOs per group, 3 separate preparations. (**e**) αSMA gene expression as assessed by qPCR after 4h of PFK15 treatment. β-actin was used as a normalization control. n=5 BVOs per group. Values are presented as mean with ± SD; P values were calculated using a two-tailed Student’s t-test. (* <0.05; **<0.005; ***<0.0005; ****<0.0001). Bar scales 50 and 100 μm.

### Proteomic analysis of the BVO secretome reveals the presence of a complex ECM structure

Maintaining microvascular integrity requires tight regulation of the EC-PC interactions. Apart from the direct contact^13^, this crosstalk also relies on the secretion of paracrine regulators^41^. BVOs are self-assembled microvascular tissue units that represent an excellent human model to study EC: PC interactions as BVOs are devoid of the major limitations of *in vitro* 2D cell culture and co-culture systems^35^. Therefore, to identify critical paracrine mediators we performed a proteomic analysis of the secretome of BVOs at baseline and following PFK15 treatment. Only proteins reported at the matrisome db plus some additional secreted proteins from our in-house generated database were considered. Using the signalP tool, 87.25% of these proteins are found to be classified as secreted based on the presence of a signal peptide (Supplementary Data Files 1 and 2).

A total of 149 proteins were identified in the conditioned media of BVOs with the top gene ontology pathway terms associated with ECM deposition (Figure 6a, Supplementary Data 1). A rich ECM comprising 58 structural and matricellular proteins and a network of 32 proteases, protease inhibitors and ECM degrading enzymes were detected. This indicated the presence of an intricate dynamic basement membrane that can adapt and respond to stimulation (Figure 6b). Transcription factor enrichment analysis revealed a regulatory network consisting of 10 transcription factors that control these differentially expressed genes (Figure 6c). The detected upregulation of the matrix-degrading enzyme matrix-metalloproteinase 9 (MMP9) in PFK15 treated BVOs (Figure 6d) was particularly interesting. Quantitative PCR analysis in BVOs revealed a sharp increase of MMP9 mRNA following PFK15 treatment (Figure 6e) and zymography confirmed enhanced enzymatic activity of gelatinases in the PFK15 treated BVO conditioned media (Figure 6f). In line with the reported inverse correlation between MMP9 and ColIV^42^, in PFK15 treated BVOs the increased MMP9 activity coincided with reduced ColIV in the basement membrane (Supplementary Figure 3a, 3b). These findings are consistent with the presence of a highly complex basement membrane in BVOs, with selected ECM structural and matricellular proteins being deposited and degraded in response to metabolic rewiring.

**Figure 6.**
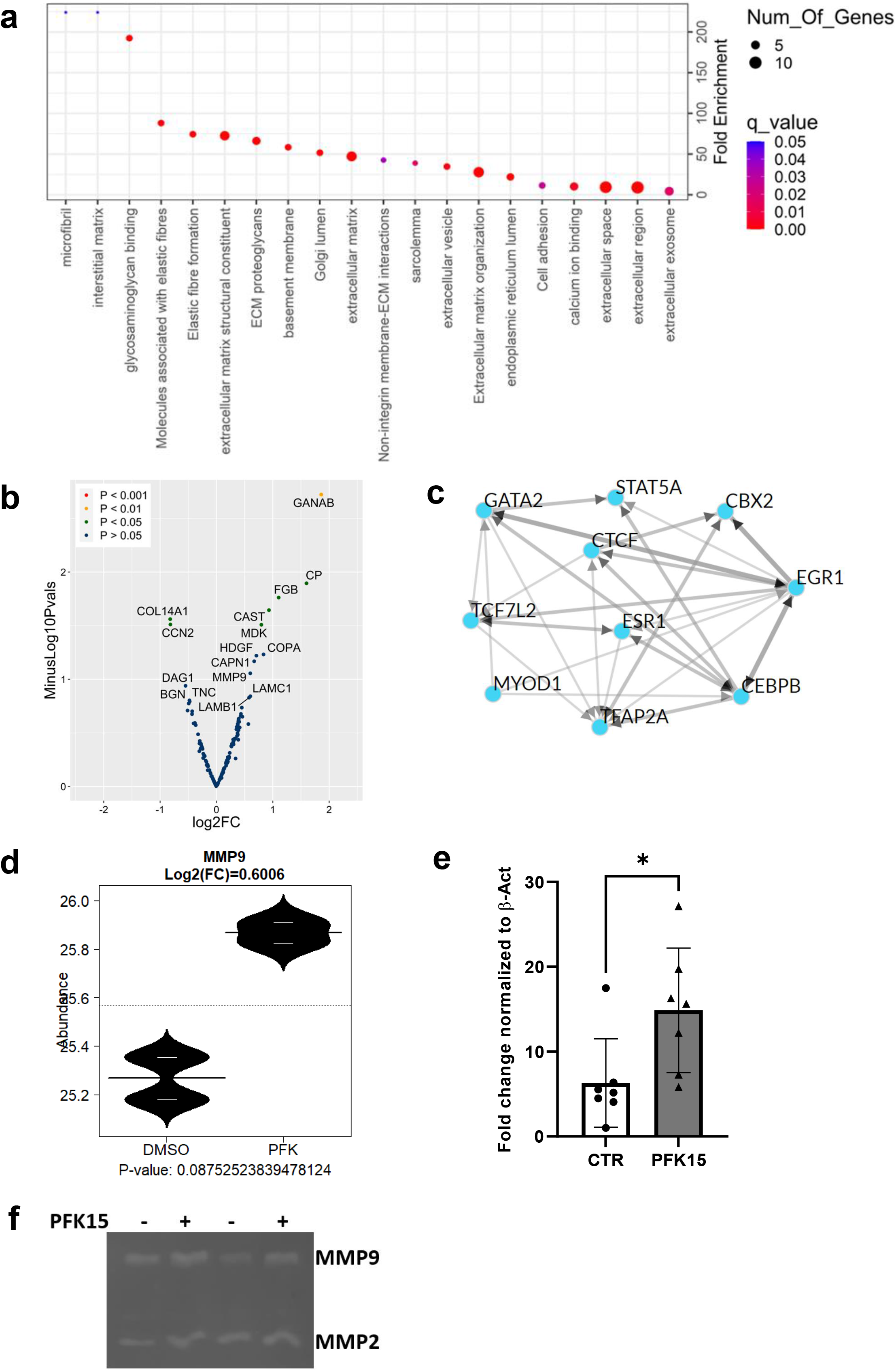
Proteomic analysis of the BVO secretome. **(a)** Pathway enrichment of proteins detected in the secretome of BVOs treated with DMSO (CTR) or PFK15 (2.5µM) for 24h. n=5 BVOs per pooled sample, from 2 separate preparations. **(b)** Volcano plot comparing the secreted proteins in conditioned media in the two groups. **(c)** Transcription factor enrichment analysis for the differentially expressed proteins using data from ENCODE database and performing analysis with the ChEA3 tool. **(d)** Differential expression of MMP9 in the BVOs secretome as detected by proteomics, n=5 BVOs per pooled sample, 2 separate preparations. **(e)** MMP9 gene expression as assessed by qPCR, β-actin was used as a normalization control. n=7 BVOs per group, 3 separate preparations, **(f)** Enzymatic activity of gelatinases in the conditioned media of BVOs treated with DMSO (CTR) or PFK15 (2.5µM) for 24h, as assessed by zymography. n=5 BVOs per pooled sample, 2 separate preparations.

### PFKFB3-driven glycolysis inhibition leads to downregulation of CTGF expression

Our proteomic analysis revealed that the cellular communication network factor 2 (CCN2, also known as CTGF), and a potent mitogen secreted by ECs^43^, were among the most downregulated proteins in the secretome of PKF15 treated BVOs (Figure 6b, 7a). In line with the proteomic data, reduced CTGF mRNA was detected by qPCR in PFK15 treated BVOs (Figure 7b) and iPS-ECs (Figure 7c).

**Figure 7.**
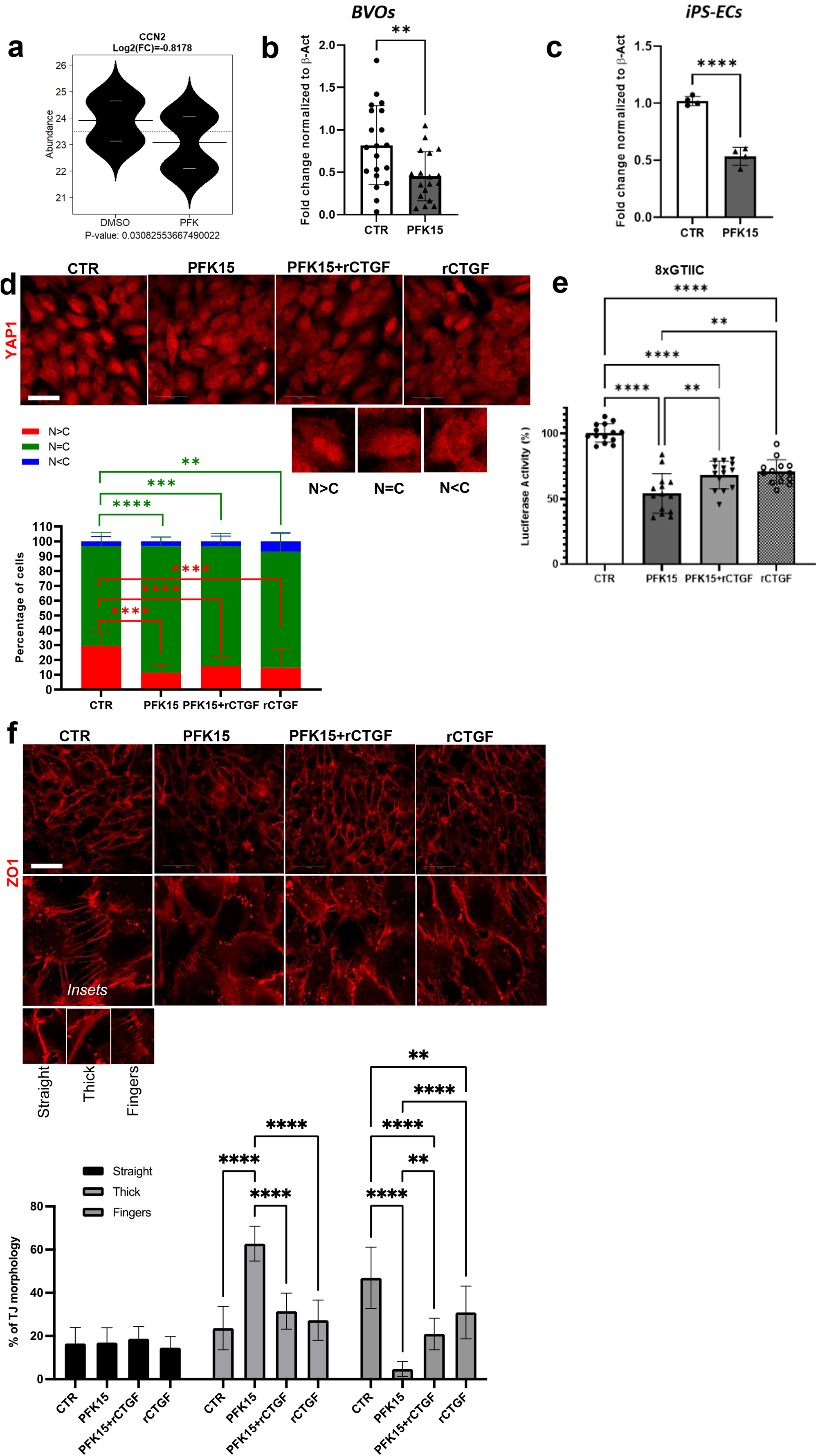
PFKFB3 mediated regulation of CTGF. The role of CTGF in iPS-EC tight junction formation. **(a)** Differential expression of CTGF in the BVOs as detected by proteomic analysis of the secretome, n=5 BVOs per pooled sample, from 2 separate preparations. **(b)** CTGF gene expression as assessed by qPCR, β-actin was used as a normalization control. n=20 BVOs per group, 3 separate preparations, **(c)** CTGF gene expression in iPS-ECs as assessed by qPCR, β-actin was used as a normalization control. n=4 independent preparations. **(d)** YAP subcellular localization following iPS-EC treatment with PFK15 for 3h, as detected using immunofluorescence confocal microscopy. Lower panel indicates quantification of YAP subcellular localization in iPS-ECs treated as indicated. N: nucleus, C: cytosol **(e)** YAP reporter activity in HEK293T cells cultured under the indicated conditions for 24h. rCTGF: recombinant CTGF. At least 3 independent transfections were assessed in quadruplicates. **(f)** Phenotypic characterization of tight junction morphology in iPS-ECs treated as indicated for 3h using immunofluorescence confocal imaging. Lower panel, quantification of straight, thick and fingers junctions. n=250 cells per group. Data are shown as mean ± SD using two-way ANOVA followed by Tukey’s multiple comparisons test. (*p<0.05; **p<0.01; ***p<0.001; ****p<0.0001)

CTGF expression is often used as a read-out of YAP activity^44^. YAP and TAZ are two transcriptional coactivators that are retained in the cytosol when rendered inactive by phosphorylation but translocate to the nucleus upon activation and bind to the TEA domain DNA-binding transcription factors initiating a cascade of events to regulate angiogenesis by controlling the EC proliferation, junction assembly and migration^45, 46^. YAP has emerged as an important mechano-transduction sensor that may also be sensitive to metabolic changes in the cells^47, 48^. In our system PFK15 treatment in iPS-ECs triggered an early reduction in the YAP nuclear fraction evident after 3h of treatment that returned to baseline after 24h (Figure 7d, Supplementary Figure 4). Similarly, YAP reporter assays indicated a decrease in YAP activity following PFK15 treatment (Figure 7e). Interestingly, recombinant CTGF supplementation could partially restore YAP activity in PFK15 treated cells, as indicated by the YAP reporter assays (Figure 7e). These findings highlight the presence of a CTGF-YAP feedforward loop in maintaining vascular integrity.

### CTGF can recover tight junction morphology

The formation of adhesion plaques and gap junctions plays a key role in the crosstalk between ECs and PCs^49^. ZO1 is a structural adaptor protein that coordinates the binding of F-actin and the transmembrane and cytosolic proteins that are required for tight junction function^45, 49^. We found that PFK15 treatment in iPS-ECs modified the junctional morphology with an increase in thick junctions, typically linked with increased permeability and a reduction in fingers junctions, which are associated with tip cells and migration^45^ (Figure 7f). Furthermore, recombinant CTGF abolished the increase in thick junctions in the PFK15 treated iPS-ECs and led to a partial recovery of fingers junctions (Figure 7f). These findings highlight the impact of PFKFB3 inhibition on YAP and its downstream effector CTGF in BVO structural remodeling and indicate an important role for CTGF in maintaining the tight junction morphology.

### CTGF supplementation recovers BVO structure following PFK15 treatment

To determine whether the PFK15-induced remodelling in BVOs can be reversed by CTGF, recombinant CTGF was used as a supplement in PFK15-treated BVOs. The self-assembled 3D vascular tissue showed again a well-organized interconnected capillary network consisting of ECs and tightly associated PCs in the control BVOs. As expected, PFK15 treatment diminished the vascular network with reduced vessel density and length accompanied by low pericyte coverage (Figure 8a, 8b, 8c). Recombinant CTGF could abrogate the PFK15 triggered structural remodelling in BVOs. BVOs treated with PFK15 in the presence of recombinant CTGF displayed vessel density and length similar to control BVOs and higher vessel density and length compared to the PFK15 BVOs (Figure 8b, 8c). This coincided with a robust and highly significant increase in PC coverage in recombinant CTGF-treated PFK15 BVOs (Figure 8d). These findings demonstrate that CTGF plays a key role in PC recruitment to capillaries and the maintenance of microvascular integrity.

**Figure 8.**
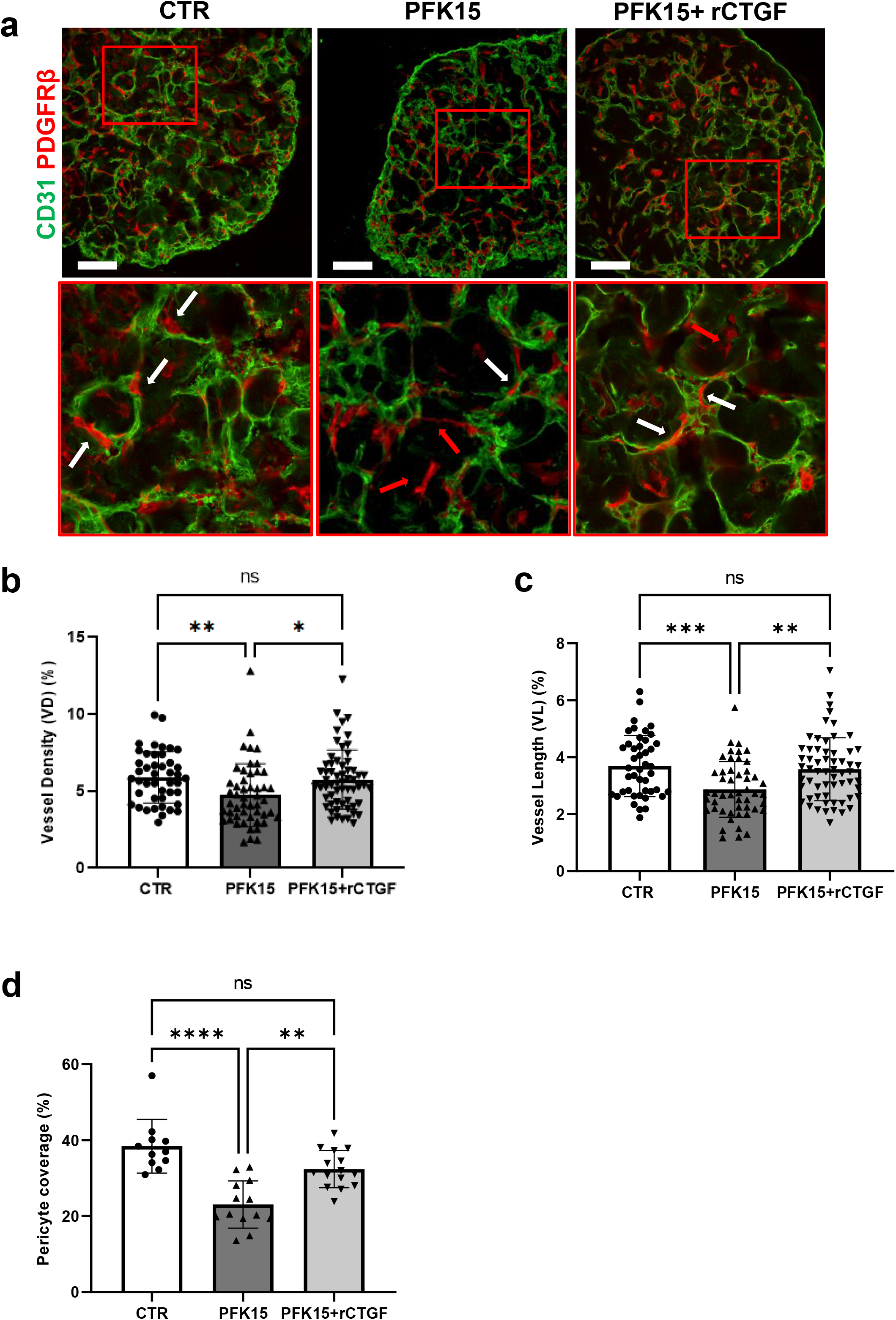
The effect of CTGF on BVO structure. **(a)** Immunofluorescence confocal imaging showing CD31^+^ ECs (green) in vascular networks covered by pericytes (PDGFRβ^+^, red) in sections from BVOs treated with DMSO (CTR) or PFK15 (2.5µM) and PFK15 (2.5µM) + rCTGF(50ng/ml) for 24h. White arrows indicate pericytes attached to microvessels, red arrows indicate ‘extravascular pericytes’. N=2 BVOs per group, from 3 separate preparations and 4 different areas per 10x images have been used for quantification **(b)** quantification of vessel density, **(c)** length and **(d)** pericyte coverage in n=2 BVOs per group from 3 separate preparations. Data are shown as mean ± SD using two-way ANOVA followed by Tukey’s multiple comparisons test. (*p<0.05; **p<0.01; ***p<0.001; ****p<0.0001). Bar scales 50 μm.

## Discussion

Here, we demonstrate that human iPS derived BVOs respond rapidly and undergo structural remodeling after short pharmacological inhibition of PFKFB3 driven glycolysis. Vessel regression, as highlighted by a reduction in vessel density and length and vessel constriction as well as by changes in microvasculature proliferation, apoptosis and transdifferentiation, was readily detectable in BVOs even after a short 24 h drug treatment. Furthermore, proteomic analysis of the BVO secretome revealed paracrine mediators implicated in ECM deposition and remodelling and identified CTGF as a critical regulator of microvascular integrity.

### Metabolic rewiring and microvascular integrity

In our experiments, PFKFB3-driven glycolysis has emerged as a key metabolic pathway in a human model of the microvasculature. Previous studies underlined the highly glycolytic nature of ECs and highlighted the advantages of using aerobic glycolysis over oxidative metabolism in the EC angiogenic potential ^15^. Here, by employing metabolic flux analysis of U-^13^C_6_ glucose, we demonstrate that iPS-ECs also rely heavily on glucose for biomass production, providing further support to the notion that human iPS-derived vascular tissue can be used as a model to faithfully recapitulate the properties and responses of the human vasculature. The metabolic rewiring induced by targeting PFKFB3-driven glycolysis in BVOs, triggered a cascade of events that had a profound effect in both the structural components of the microvasculature, namely ECs, PCs, basement membrane- and their interactions.

Similar to the PFKFB3 deletion in ECs in mice^15^, we observed inhibition of EC proliferation, increased quiescence and impaired expansion of vascular plexus in PFK15 treated BVOs. A significant decrease in vessel density and length was noted following glycolysis inhibition in BVOs. In contrast to ECs, PFKFB3 inhibition did not affect PC proliferation or apoptosis but instead affected their location and differentiation state. In terms of cell localisation, we detected a pronounced PC dropout (∼50%) in human PFK15 treated BVOs, contrary to the findings in mice harbouring a deletion of PFKFB3 in ECs, where no changes were detected ^15^. Although the mechanism is currently unclear, the differences in the timeframe of PFKFB3 inhibition (chronic in PFKFB3 mutant mice versus short-term in BVOs) and interspecies differences (mouse versus human) could explain the different phenotypes.

The tight interactions between ECs and PCs are critical for the integrity of mature capillary networks and the detachment of PCs from the microvasculature has a destabilizing effect on microvessels^50^. Noteworthy, pioneering studies in PDGFβ+/- mice demonstrated that a 50% reduction in PC density can have a causal effect in triggering pathological angiogenesis^9^. In PFK15 treated BVOs no difference in PDGFRβ expression level was observed (data not shown), but the 50% decline in PC coverage was accompanied by microvascular regression with reduced vessel density and length. Even though the sequence of these events is not clear, the proteolytic degradation of the basement membrane by MMPs has been implicated in the detachment of PCs^51^. Hence, the increased presence and proteolytic activity of MMP9 in the PFK15 BVOs as confirmed by proteomic analysis and zymography may facilitate the increased release of PCs from the basement membrane, a concept that is also supported by the reduced thickness of ColIV in the PFK15 treated microvessels.

Following the drop in microvessel PC coverage, the extravascular PCs differentiated into SMC-like cells in PFK15 treated BVOs. A pronounced increase in αSMA expression and to a lesser extent an upregulation of SM22 were detected. This unexpected finding may indicate activation of PCs in PFK15 treated BVOs. Several studies have demonstrated that PCs display high phenotypic heterogeneity ^13^. For example, these cells can be αSMA^+^ as is the case for PCs in post-capillary venules or αSMA^-^ PCs along microcapillaries *in vivo* ^52^. PCs that are cultured *in vitro* or are specifically activated can express αSMA ^53^. Previous reports have identified opposing actions of TGFβ1 and FGF2/ IL-6 in PC differentiation. In a mouse model of retinal angiogenesis, FGF2 and IL-6 were shown to regulate PC coverage while TGFβ1 signalling promoted human pulmonary PC differentiation into contractile SMC-like cells *in vitro*^*39, 40*^. Intriguingly, increased protease activity leading to proteolytic degradation of the ECM can also release growth factors such as FGF2 and TGFβ1 and actively contribute to changes in the cellular microenvironment that may affect the PC differentiation state^42^. Thus, there is a delicate balance between negative and positive regulators of PC differentiation that determines the activation state and the contractile function of PCs in the microvasculature^54^. In our system, the impact of PC transdifferentiation to SMC-like cells is not clear. PCs triggered mechanical modifications of their microenvironment that directly transmit mechanical strain to ECs through the underlying substrate has been proposed^55^, but further studies are required to test and confirm this hypothesis.

### CTGF and microvascular integrity

CTGF is a key regulator of ECM deposition with distinct roles in development and in disease^56^. CTGF null mice die shortly after birth due to respiratory defects^56^, but studies in developing CTGF mutant embryos have revealed an important role of CTGF in PC recruitment to the vasculature^57^. Noteworthy, reduced deposition of ColIV was also observed in CTGF mutants, thus indicating a role for CTGF in basement membrane formation^57^. Similarly, in a mouse model targeting CTGF expression specifically in ECs^58^, impaired vascular cell growth and morphogenesis was detected in the postnatal retina. In these mice a key role for CTGF-YAP in retinal vascular development was reported^58^. In disease, in a mouse model of diabetic retinopathy, CTGF expression was required for thickening of the basal lamina of retinal capillaries, an instrumental step in the development of vision loss^43^ while in diabetic nephropathy, the increased CTGF expression was shown to inhibit BMP-7 signal transduction, reduce MMP activity and contribute to basement membrane thickening and albuminuria^59^.

In patients, targeting CTGF has mainly been associated with the development of fibrosis. CTGF was proposed as a surrogate marker of fibroproliferative disease for several pathologies^60^ while another study has reported preferential accumulation of the NH2-terminal CTGF fragment in proliferative diabetic retinopathy^61^. By far the most encouraging results were obtained in a phase 2 clinical trial using pamrevlumab (FG-3019), a recombinant human monoclonal antibody against CTGF, that was shown to be safe and effective in attenuating the progression of idiopathic pulmonary fibrosis^62^.

In our organoid model of human microvasculature, CTGF emerged as a paracrine regulator that can alleviate the vascular remodelling induced by the metabolic rewiring, by reinforcing the direct interaction of ECs and PCs and recovering the morphology of tight junctions. Our study highlights the presence of a CTGF-YAP feedforward loop in maintaining vascular integrity. We provide evidence that PFKFB3 is an upstream regulator of YAP, affecting its nuclear localization and transcriptional activity. Supplementation with recombinant CTGF can partially restore YAP activity, recover the alterations in tight junction morphology induced by inhibition of glycolysis, and effectively correct the ‘leaky’ thick junctions bringing them to the levels observed in the control BVOs, enriching tight junctions for the ‘migratory’ fingers junctions. This normalization in tight junction morphology has a beneficial effect on EC: PC direct interaction in BVOs. In our 3D model of the microvasculature, we observed a significant increase in PC coverage in microvessels and vessel density and length, indicating that CTGF can protect the microvasculature from structural remodelling triggered by metabolic rewiring.

## Conclusions

Our study indicates that BVOs respond rapidly to metabolic changes. The prompt structural remodelling that is observed, makes this model extremely attractive for mechanistic studies and the identification of potential targets for novel therapeutic approaches. Importantly, this short timeframe of response is also useful and desirable for high throughput drug and small compound screening applications. Furthermore, the use of patient derived iPS cells and BVOs that can capture the interpatient variation and predict drug responses more accurately may facilitate the development of personalized treatment strategies.

## Acknowledgements

This study was supported by King’s BHF Centre of Research Excellence (RE/18/2/34213), the British Heart Foundation (PG/19/56/34550) and the European Research Council (ERC) Advanced Grant 787971 “CuRE”.

## Author contributions

SR and AZ conceived the idea and designed the experiments. SR, IS, ES, CR, CS, AP, XY and AZ performed experiments. KT performed the bioinformatic analysis of the proteomic dataset. All the authors participated in data collection and interpretation. AZ drafted the manuscript with critical revision feedback from all other authors. All authors read and approved the final manuscript.

---

**Supplementary Movie 1**

Confocal stack of a CTR BVO stained for CD31 (green), PDGFR-β (red) and DAPI (blue).

**Supplementary Movie 2**

Confocal stack of a PFK15 treated BVO stained for CD31 (green), PDGFR-β (red) and DAPI (blue).

## Supplementary Figures

**Supplementary Figure 1.**
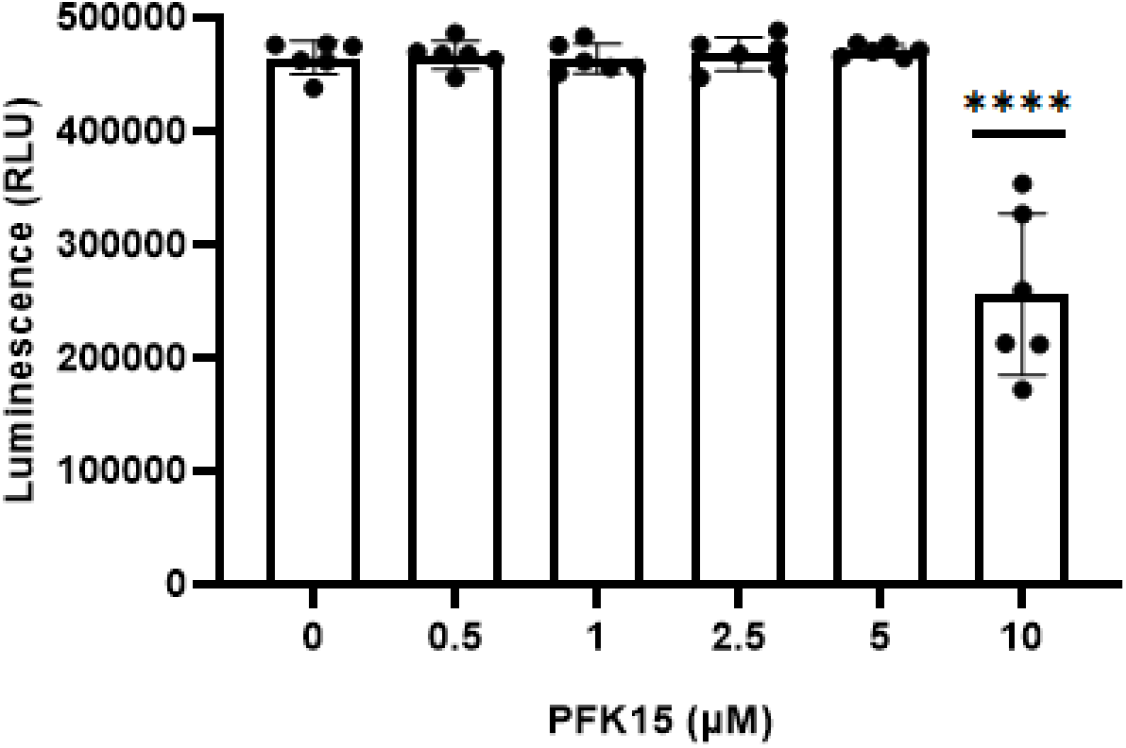
Survival of iPS-ECs treated with various concentrations of PFK15 for 24h. ****p<0.0001

**Supplementary Figure 2.**
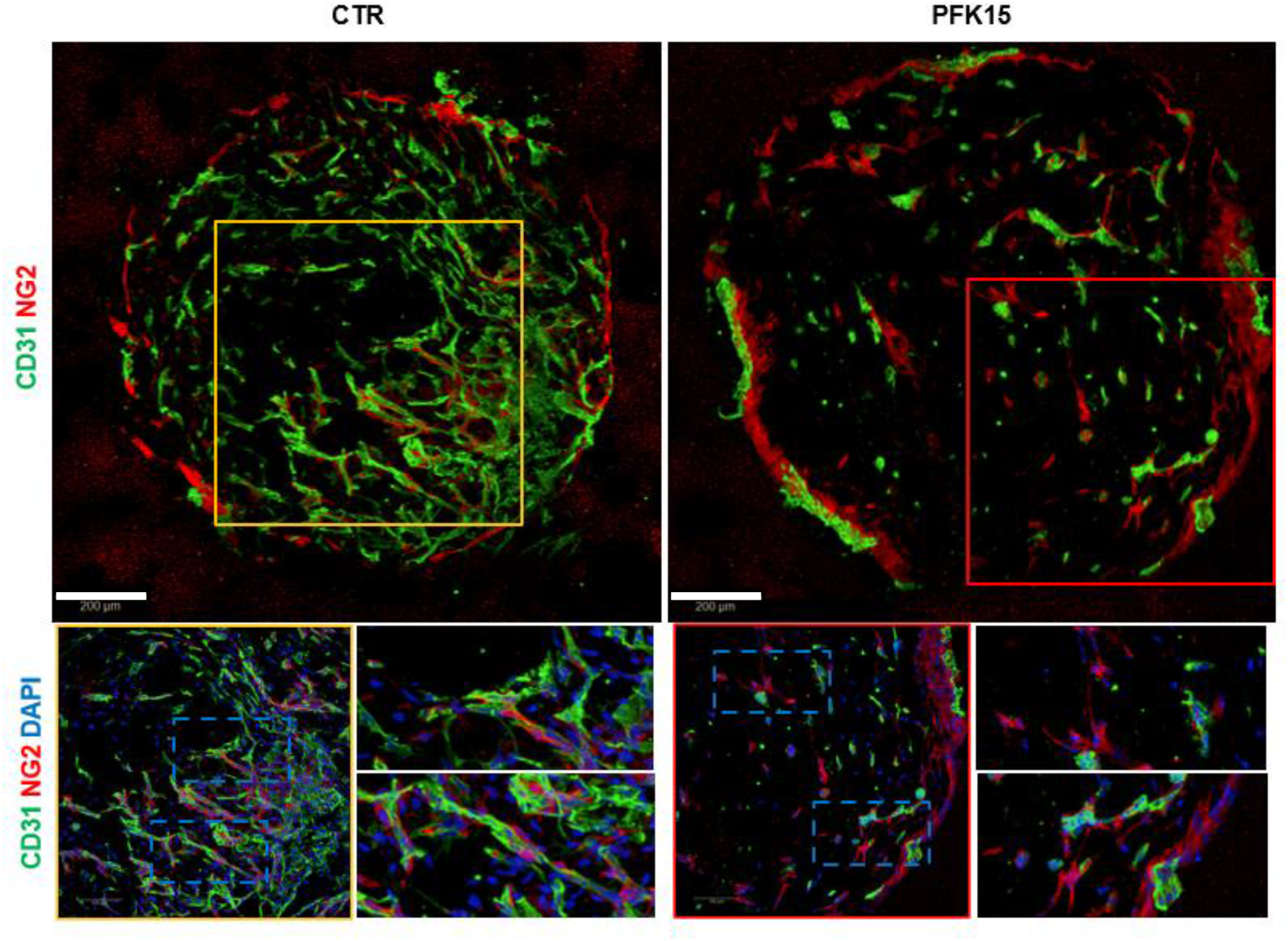
NG2 expression in BVOs following PFK15 treatment. Phenotypic characterization of NG2 (red) expression in BVO sections using immunofluorescence confocal microscopy. CD31^+^ ECs are shown in green. Bar scales 200 μm.

**Supplementary Figure 3.**
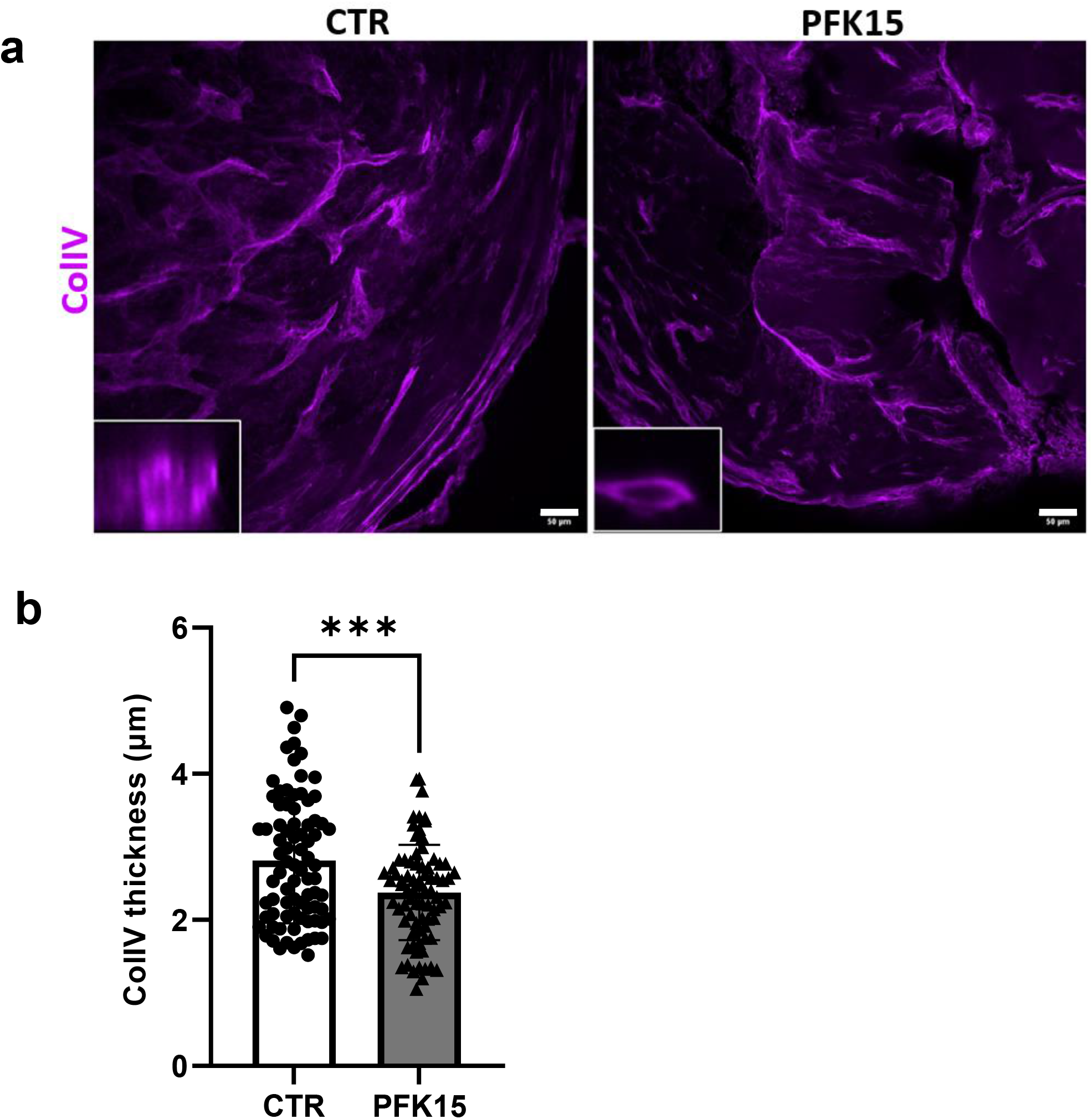
Deposition of ColIV in BVOs following PFK15 treatment. **(a)** Representative images of basement membrane, as detected by immunofluorescence confocal imaging for ColIV in sections from BVOs treated with DMSO (CTR) or PFK15 (2.5µM) 24h. N=80 cross-sections in single BVOs from 2 separate preparations per group **(b)** Vessel cross-sections were used to quantify basement membrane thickening. Values are presented as mean with SD; P values were calculated using a two-tailed Student’s t-test. (***<0.0005). Bar scales 50 μm.

**Supplementary Figure 4.**
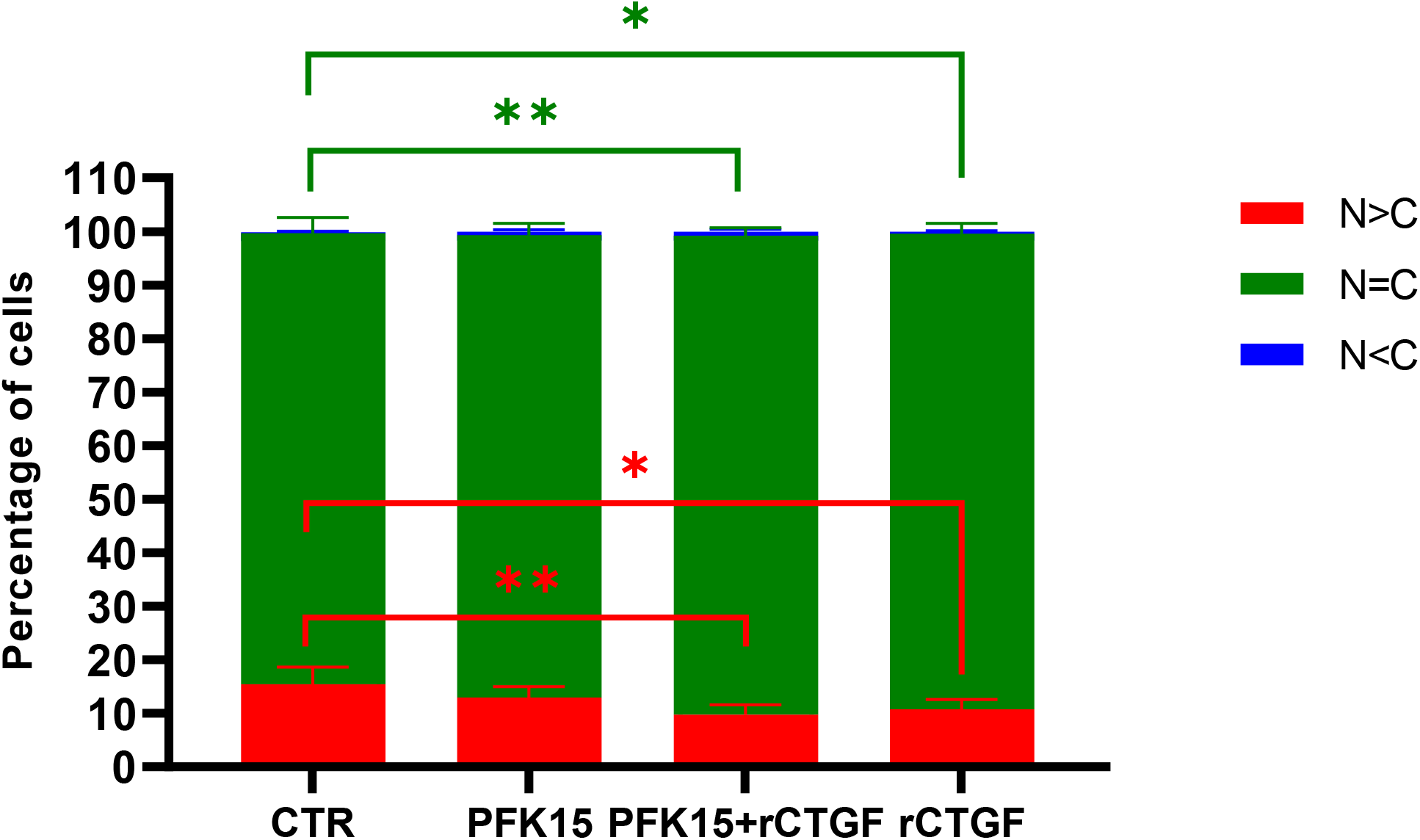
YAP subcellular localization in iPS-EC with the indicated treatments for 24h as detected using immunofluorescence confocal microscopy. Data are shown mean with SD using two-way ANOVA followed by Tuckey’s multiple comparisons test. (*p<0.05; **p<0.01).

