## Supplementary Info for "Human blood vessel organoids reveal a critical role for CTGF in maintaining microvascular integrity"

Sara G Romeo^1^, Ilaria Secco^1^, Edoardo Schneider^1^, Christina M Reumiller^1^, Celio XC Santos^1^, Aman Pooni^1^, Xiaoke Yin^1^, Konstantinos Theofilatos^1^, Silvia Cellone Trevelin^1^, Lingfang Zeng^1^, Giovanni E Mann^1^, Andriana Margariti^2^, Manuel Mayr^1^, Ajay M Shah^1^, Mauro Giacca^1^ and Anna Zampetaki^1*^

^1^King's College London British Heart Foundation Centre, School of Cardiovascular Medicine & Metabolic and Sciences, London, United Kingdom; ^2^ The Wellcome-Wolfson Institute for Experimental Medicine, Queen’s University Belfast, Belfast, United Kingdom.

**Supplementary Material**

**Supplementary Table S1**

Antibodies used for Immunofluorescence staining

| **Primary antibody** | **Company Reference** | **Dilution IMF** |
| --- | --- | --- |
| CD31 | R&D System AF806 | 1:100 |
| PDGFRβ | Cell Signaling #3169 | 1:100 |
| Collagen IV | Millipore AB769 | 1:200 |
| Oct4 | Thermo TA500035 | 1:100 |
| Nanog | Sigma N3038 | 1:100 |
| CD144 (VE-cadherin) | Millipore MABT134 | 1:200 |
| ZO1 | Santacruz Technologies sc-8147 | 1:100 |
| αSMA | Sigma C6198 | 1:300 |
| SM22 | Abcam ab14106 | 1:500 |
| YAP1 | NOVUS NB110-58358 | 1:500 |
| NG2 | Abcam ab86067 | 1:100 |
| Ki67 | Cell Signaling #9129S | 1:100 |
| CC3 (Cleaved-Caspase 3) | Cell Signaling #9661S | 1:250 |

| **Secondary antibody** | **Company Reference** | **Dilution IMF** |
| --- | --- | --- |
| Alexa-Fluor 488 Donkey anti-Sheep | Invitrogen A11015 | 1:250 |
| Alexa-Fluor 488 Donkey anti-Mouse | Invitrogen A21202 | 1:250 |
| Alexa-Fluor 488 Donkey anti-Rabbit | Invitrogen A21206 | 1:250 |
| Alexa-Fluor 555 Donkey anti-Rabbit | Invitrogen A31572 | 1:250 |
| Alexa-Fluor 633 Donkey anti-Mouse | Invitrogen A21100 | 1:250 |
| Alexa-Fluor 647 Donkey anti-Rabbit | Invitrogen A31573 | 1:250 |
| Alexa-Fluor 647 Donkey anti-Goat | Jackson Immunolabs 705-606-147 | 1:250 |

| **Gene** | **Forward Primer** | **Reverse Primer** |
| --- | --- | --- |
| **αSMA** | CCTGACTGAGCGTGGCTATT | GCCCATCAGGCAACTCGTAA |
| **MMP9** | CCTGGGCAGATTCCAAACCT | CAAAGGCGTCGTCAATCACC |
| **CCN2 (CTGF)** | CCGCACAAGGGCCTATTCT | GGTACACCGTACCACCGAAG |
| **B-Actin** | CGTCTTCCCCTCCATCGTG | CTCGATGGGGTACTTCAGGG |

**Supplementary Table S2**

List of Primers used for quantification of gene expression

**Supplementary Table S3**

List of antibodies used for FACS Analysis

| **Primary antibody** | **Company Reference** | **Dilution** |
| --- | --- | --- |
| CD31-AlexaFluor647 | BD Biosciences, 558094 | 5µl/test |
| CD140b-PE | BD Biosciences, 558821 | 20µl/test |
| CD144-FITC | BD Biosciences, 560874 | 20µl/test |
| CD45-FITC | Invitrogen, 11-0459-41 | 5µl(0.25µg)/test |
| CD90-PerCP/Cyanine5.5 | Biolegend, 328117 | 1µg/million cells |
| CD73-BV650 | BD Biosciences, 742633 | 10µl/test |
| CD44-PE | BD Biosciences, 550989 | 20µl/test |
| CD144-BV786 | BD Biosciences, 565672 | 5µl(0.25µg)/test |
| Live/Dead-FVS780 | BD Biosciences, 565388 | 0.1µl/test |

**Supplementary Table S4**

List of antibodies used for Western blot Analysis

| **Primary antibody** | **Company Reference** | **Dilution WB** |
| --- | --- | --- |
| CD31 | Abcam ab28364 | 1:500 |
| CD144 (VE-cadherin) | Millipore MABT134 | 1:2000 |
| KDR (VEGFR2) | Cell Signaling #2479 | 1:1000 |
| eNOS | BD Biosciences 610297 | 1:5000 |
| Oct4 | Thermo TA500035 | 1:1000 |
| GAPDH | Santacruz Technologies sc-25778 | 1:1000 |

| **Secondary antibody** | **Company Reference** | **Dilution WB** |
| --- | --- | --- |
| Peroxidase AffiniPure Goat Anti-Mouse IgG | Jackson Immunolabs 115-035-174 | 1:4000 |
| Peroxidase IgG Fraction Monoclonal Mouse Anti-Rabbit IgG | Jackson Immunolabs 211-032-171 | 1:4000 |

**Supplementary Table S5**

| **P value** | **Corrected P-value (q value)** |
| --- | --- |
| 0.0002 | 0.0012 |
| 0.0237 | 0.0474 |
| 0.0006 | 0.0018 |
| 0.5450 | 0.5450 |
| 0.2065 | 0.2478 |
| 0.0725 | 0.1088 |
