## Supplementary Movies for "Human blood vessel organoids reveal a critical role for CTGF in maintaining microvascular integrity"

### Slide 1
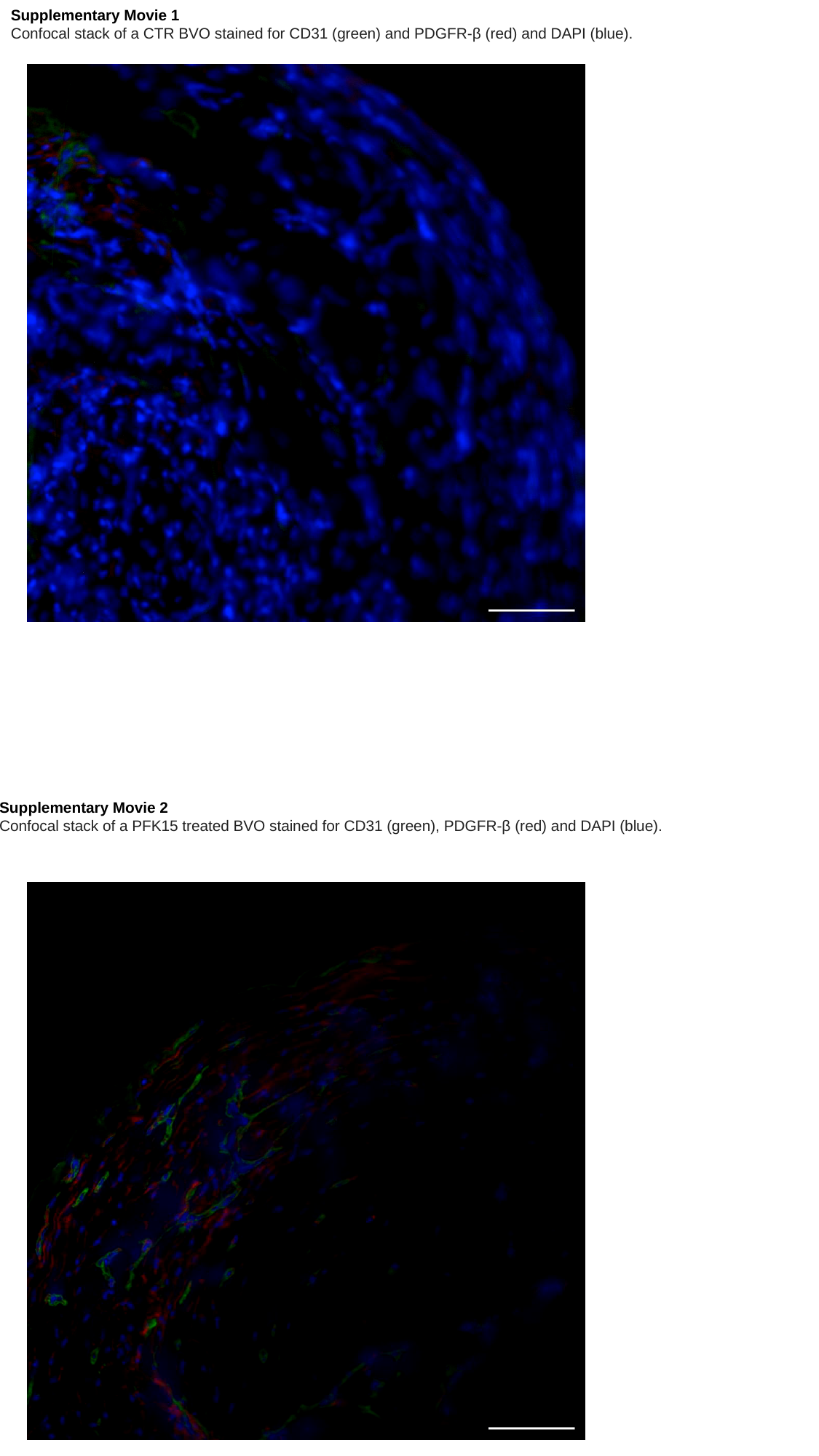

Supplementary Movie 1
Confocal stack of a CTR BVO stained for CD31 (green) and PDGFR-β (red) and DAPI (blue).
Supplementary Movie 2
Confocal stack of a PFK15 treated BVO stained for CD31 (green), PDGFR-β (red) and DAPI (blue).
